# Gait analysis of Pak Biawak: a necrobot lizard built using the skeleton of an Asian water monitor (*Varanus salvator*)

**DOI:** 10.1101/2025.07.12.664518

**Authors:** Leo Foulds, Donan Satria Yudha, Parvez Alam

## Abstract

In this paper,we consider the feasibility of mimicking the sprawling gait of a live varanid (*Varanus salvator*) using a necrobot (named: Pak Biawak), a robot constructed using the skeletal parts of a deceased varanid of the same species. Pak Biawak is manufactured using simple joints and components, and limb motion is coupled to passive spine bending to enable the sprawling gait. Here, we assess both the lateral and dorsal kinematics of Pak Biawak at different speeds, and compare the metrics from each to those of a similarly sized live varanid. When assessing lateral view shape metrics (stride aspect ratio, stride circularity, normalised stride swept area, normalised stride swept area perimeter), we find that Pak Biawak’s gait is consistent across all speeds and the majority of Pak Biawak’s lateral shape metrics are kinematically aligned with those of the live varanid. This also proves true when comparing Pak Biawak’s lateral trajectory metrics (radial distance of swept area, normallised root mean squared error) against those of the live varanid, and at different speeds of sprawling. Pak Biawak’s dorsal metrics include the spine bending amplitude and period, and these are not found to be significantly different to those of the live varanid, however, Pak Biawak’s amplitude is affected by sprawling speed. We use three metrics to compare forward and reverse limb sweeps including, angular curvature, differential curvature, and a normalised arc length. Of these, a preponderance of highly significant differences (p *≤* 0.001) are observed on comparing the forward sweep arc length of Pak Biawak at every sprawling speed against the forward sweep arc length of the live lizard. All other kinematic metrics in the necrobot are nevertheless very close to those of the live lizard. Finally, when comparing the trackway width of Pak Biawak against the live lizard, we again find there is very close kinematic compatibility between the two, and conclude that our necrobot can be designed and manufactured to mimic the sprawling gait of a real varanid, even when using simple kinematic linkages in unison with a passive spine bending differential applied at only one central location in the necrobot spine.

## 1. Introduction

Lizards belong to the Squamata clade, which are believed to have evolved approximately 150 million years ago [1]. They are found across a broad range of different habitats, from semi-aquatic to deserts, to jungles and forests, exhibiting the capacity to walk, run, [2,3] swim [4], climb [5] and stand upright [6]. The study of their gait [2,3,7–9] has provided several insights on adaptable, efficient movement, which is particularly relevant in the field of mobile bio-inspired robots. While lizards are tetrapods, their walking posture is not typically upright, such as can be observed in mammals like dogs and horses [10]. Lizard locomotion is often described as sprawling, where the limbs protrude laterally from the body and move in a sweeping motion [11]. Lizard gait generates a lateral undulation of the spine, which sometimes changes from a standing wave in slower gaits to a travelling wave in faster gaits [12][13]. The lizard sprawling gait allows for a low centre of gravity [12], which might aid stability, whilst concurrently improving climbing ability [14]. In fact, climbing species of lizards should can adjust their gait to counter caudally directed (when climbing up) or cranially directed (when climbing down) gravitational forces [15]. Some species of lizard, for example, the *Varanus salvator*, or the water monitor, is a predominantly land-dwelling lizard possessing a strong ability to swim [1].

The typical lizard gait has numerous locomotive benefits that could be applied to legged mobile robot locomotion. Specifically, the sprawling mode of locomotion maintains a low centre of gravity whilst also enabling multiple points of contact with the ground, including from the tail. This stance may have the benefit of providing additional stability in a mobile robot on challenging terrain. Sprawling locomotion nevertheless, is not ideal for all terrain and problems may arise in tight spaces due to its wide lateral motion. Nevertheless, spinal undulation, when coupled to sprawling movement, might decrease the overall space restrictions for this gait. Additionally, the natural flexibility of a segmented body has the potential to improve a robot’s ability to navigate obstacles such as this [16]. Such adjustments can be observed in insects such as ants and cockroaches, where gait adjustments improve stability when bearing load or changing speed [17,18]. These types of adjustments provide the basis for various robot designs [19,20], an example of which is the robot Ajax, who incorporates 24 degrees of freedom to mimic the agility and speed of a cockroach. Other less typical adjustments include rolling [21], as observed in the huntsman spider, who will rather roll than walk to enable faster and more efficient travel, a coupled kinematic mimicked in the quadruped robot, Bilbiq [22]. Recent examples of sprawling in the design of robots include RSTAR [23], a simple structured sprawling robot who could overcome challenging obstacles including compliant as well as slippery surfaces, and who could climb vertically in a tube or between two walls. RSTAR had essentially two limbs, which is a very different construction to the typical four limbed sprawl observed in lizards. A four legged example of a robot sprawler is TITAN-XIII [24], who has a complex appendage system to enable 3 degrees of freedom (DoF) of movement to enable the sprawling gait. Sprawling has been designed into wheeled robots like the Passively Sprawling Robot (PSR) [25], whose sprawl capabilities are from the inclusion of a lateral spring mechanism that adjusts passively with the turning of its legs, which are themselves designed into circular wheel-like rotational structures. In bioinspired robots that aim to mimic the structure and shape of real lizards, there is importance in enabling spine movement, as noted by Horvat et al. [26], who designed robotic sprawling using a side-to-side spine controlling mechanism, specifically one which is independently controlled with sufficient DoF at both the shoulder and pelvic girdles. There is also importance in considering tail improved stability, as evidenced through Tail STAR [27], a wheeled sprawling robot that used a bi-segmented tail to improve Tail STAR’s ability to climb high staircases. While robotic sprawlers can benefit from the incorporation of complex appendage designs [28,29], simplifying appendage designs can still result in clear kinematic benefits [29]. The design of robotic structures with kinematic simplicity is of considerable importance, since it can benefit control accuracy by reducing the complexity of the mathematical models needed to describe and control robot motion, minimises potential positioning errors through the minimalistic utility of links and joints, decreases actuator count and hence energy input requirements, and decreases material use and waste [30–33].

While bioinspired robots take many shapes and forms, they are by and large predominantly manufactured using processed plastics and metals. It is expected that by 2050 there will be 12 billion tonnes of plastic waste in the environment [34]. The processed materials commonly used in robotics are seen as beneficial due to their weight, durability and properties [35]. These materials are nevertheless, extremely resistant to biodegradation [36] with less than 10% of all plastics being commonly recycled [37]. Robots are frequently utilised in nature. COTSbot for example, is a robot used to eliminate invasive starfish species [38]. While it has an obvious ecologically friendly intention, the robot is made from plastic, potentially introducing microplastics, such as polypropylene and other polymers, into the ecosystem [39]. Biomimetic soft robots [40–43] and composite robots [44–47] are also gaining popularity and it is expected that they will be used in natural environments as soft robotics technologies mature [48–51]. Many soft robots are made using elastomeric materials such as silicone [52], leachates from which are known to be detrimental to the marine environment. The introduction of existing biological materials into robot design is a means by which the use of plastic in bio-inspired legged mobile robots could be reduced.

Though there are increasing efforts to use natural, and ecologically more friendly materials in robotics [53–57], there are still considerable environmental burdens from these materials, from harvesting to processing and waste disposal [58–60]. Necrobots may solve some of these environmental problems. A necrobot is a robot that repurposes post-mortem body parts, taking advantage thus, of naturally manufactured functional structures. Animal parts, such as bones and exoskeletons, are ubiquitously available and are biodegradable, while concurrently being rigid, lightweight structures that can bear load and be re-engineered to move. As such, they have the potential to replace the synthetic materials currently used in robotics. While bones do often exhibit a toughness reduction and stiffness increase after death, due to the degradation of collagen and dehydration [61], they can be easily and inexpensively replaced. Animal bones make up 100 million tonnes of waste due to the consumption of meat products [62], bones that could easily be re-purposed to form necrobots with potential benefits to society. They provide an alternative to using non-biodegradable materials such as plastic or metal and require little to no subtractive manufacturing as they are pre-formed and have already been field tested in nature. There are only a few necrobot studies to date, but each has shown considerable potential. The spider necrobot end effectors developed by Yap and co-workers [63] have been shown to carry 130% their own body mass, while carefully designed metastructures integrated as artificial joints within the necrobot crow ‘Sundoli’ enable a payload ratio of 1400% [64]. In fact, the highest recorded payload ratio of any walking robot belongs to ‘Poka’, an ambulating necrobot beetle developed by Tsvetkov and Alam [65], with a measured payload ratio of 6847%.

In this paper, we consider the design, manufacture and testing of a ca. 2 m long varanid necrobot, hereafter named, ‘Pak Biawak’. Rather than focus on payload capacity or transport cost, we focus on mimicking and characterising the lizard sprawling gait using a varanid endoskeleton as the main body of the robot. In this paper, we compare Pak Baiwak’s gait against the gait of a real varanid lizard using a range of lateral view and dorsal view kinematic metrics. The research reported herein represents the first full varanid skeleton repurposed to form a sprawling necrobot.

## 2. Results and Discussion

### 2.1. Design and manufacture

A series of different mechanical parts were designed to enable connectivity and motion initially in a prototype robot, additively manufactured using the 3D scanned endoskeletal segments of a real varanid. The 3D printed prototype was used to enable the judicious design and testing of parts and components, as well as to aid in determining key locations for the final necrobot, without damaging the real skeleton. Once the parts and positioning was ascertained, the parts and components were connected to the necrobot, Pak Biawak, using the original skeleton from which the 3D scans were made. The parts/components included specialised spine clips, which were designed as connections to the spine, Figure 1(a), a differential gearbox to allow for lateral (from −90*^◦^* to +90*^◦^*) and dorsoventral spine bending between 0*^◦^* and 90*^◦^*, Figure 1(b), keys to transfer bending between the differential gearbox and spine clips, Figures 1(c-d), a servo mount bracket shown to fix the motors to the spine using the spine clips, Figure 1(e), a rotary bar to both fix the servo motor while providing a 6 cm diameter step circle, Figure 1(f), a rotary fork interfaced to the skeleton limbs and attached to a linkage using a press fit bearing while the two arms were connected to the limbs using PolyMorph plastic, Figure 1(g), support bars, Figure 1(h), to limit motion to 1 DoF in order to favour a stable gait, bolt locator brackets, Figure 1(i), to support the walking mechanism, and an electronics box to house and protect the electronics, Figure 1(j), which includes numerous slits to enable the convenient routing of cables.

**Figure 1.:**
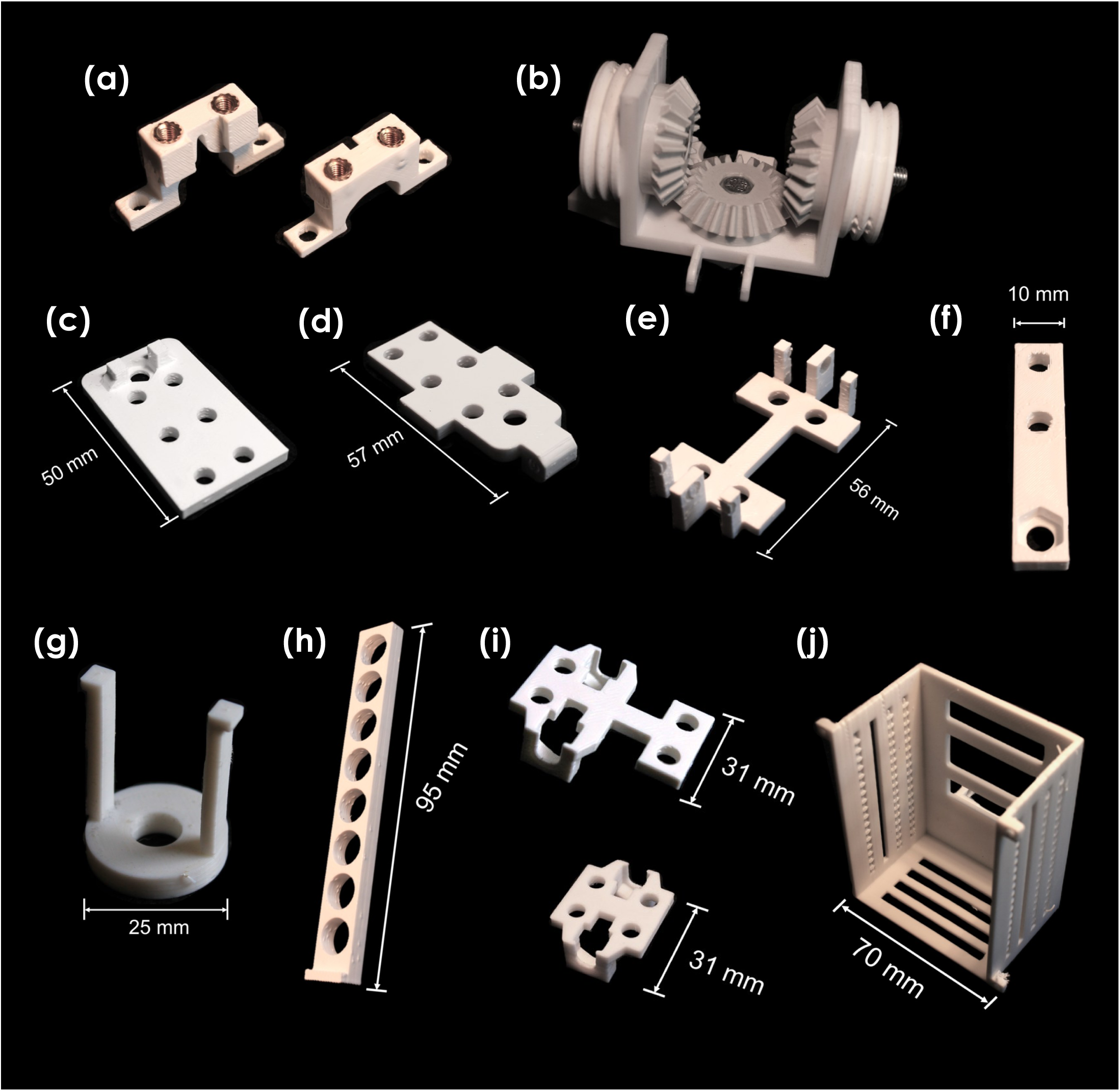
Parts designed and additively manufactured to enable connectivity and motion in the robot: (a) spine clips (b) differential gearbox (c-d) keys (e) servo mount bracket (f) rotary bar (g) rotary fork (h) support bars (i) bolt locator brackets (j) electronics box.

The differential gearbox was affixed to the centre of the spine, Figure 2(a). One key was pinned to allow rotation between two arms protruding from the housing, which facilitated dorsoventral bending. Another key was fixed to the bottom bevel gear to rotate in the horizontal plane, facilitating lateral bending. Two bolt locators were secured beneath the keys to form part of the walking mechanism. Bolts were secured through both the keys and bolt locators onto the spine clips. The walking mechanism was based on a four-bar linkage mechanism, specifically the crank-rocker variation. The basic crank-rocker mechanism is shown in Figure 2(b). Full rotation occurs at point 1, where the crank drives the orange bar in a circular motion. At point 2, a passive joint allows full rotational movement, creating a rocking motion at points 3 and 4. The linkage at point 3 stabilises the walking movement, creating an opposing force when there is resistance at the end of the blue bar. The blue bar represents the skeleton limb, while the grey bar represents the spine. Four servo motors were mounted onto the spine with rotational shafts facing outwards. The humeri and femora were mounted onto the servo motors via the forks (see Figure 2(c) and (d)) such that they could rotate on the rotating shaft. Support bars were mounted to the elbow and knee joints of each limb using mouldable PolyMorph. The space between the support bars was bridged by an M5 bolt. The base of the rocking bar (i.e., the green bar in Figure 2) was attached to the bolts protruding from the bolt locators.

**Figure 2.:**
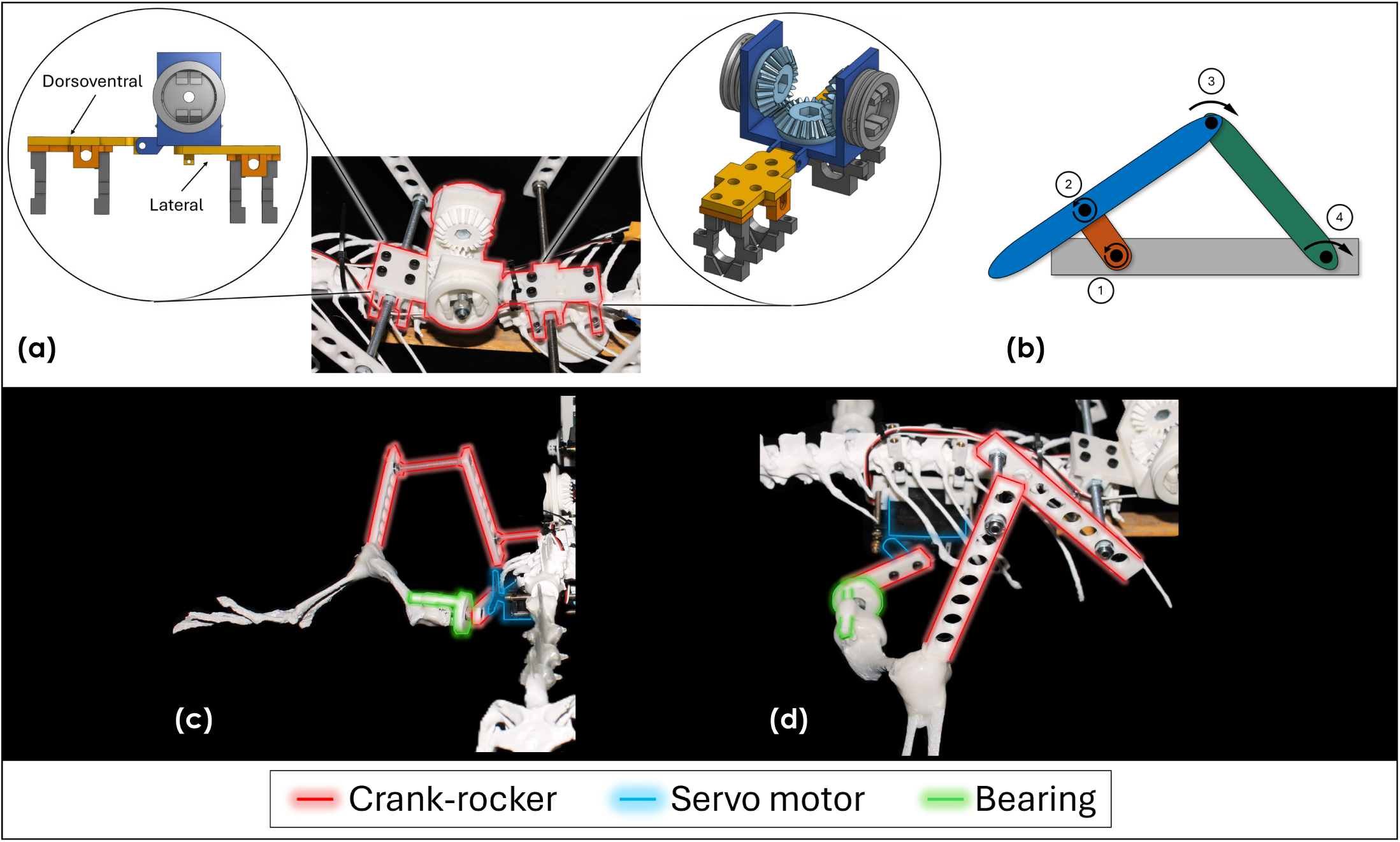
(a) differential gear box for spine bending fitted with clips for spine connection (b) crank-rocker mechanism for walking shown from the anterior view, and crank-rocker mechanism shown from (c) anterior and (d) lateral views.

The servo motors were sized to fit within the varanid skeleton ribcage. Considering the torque and dimension requirements, we selected a Parallax Inc 4 servo motors (42.5g). This motor features 27Ncm (0.9kg at 3cm) of torque at 6V, which provides a margin of error of approximately four for each one. The servo motors featured pulse width modulation (PWM) control for rotational speed (up to 50 RPM), so ambulation velocities could be varied during experimentation. Each servo required 4-6V, having a maximum current draw of 140±50mA when rotating with no load. The static current draw was 15mA. As a safety buffer, it was assumed that each servo would draw 1A in the worst-case scenario, such that a minimum of 4A would be supplied to the necrobot.

An Arduino Uno R3 was used to control the servo motors via the PWM pins to adjust the direction and speed of rotation. The datasheet for the R3 [66] defines signal pulses of 1.3 ms for full clockwise speed, 1.5 ms for stationary, and 1.7 ms for full anticlockwise speed. To mimic the sprawling gait of the varanid *in-vivo*, opposing fore and hind limbs were moved in sync. Clockwise rotation moved the hind limbs forward, while anticlockwise moved the forelimbs forward. One forward step consisted of a clockwise (*<*1.5 ms) and anticlockwise (*>*1.5 ms) rotational pair, with pulse lengths varied in 25 *µs* steps. To simplify the tested speeds, the naming convention displayed in Table 1 will be followed hereafter.

**Table 1.:** PWM pulse length settings (in *µs*) for approximated rotational speeds.

|  | PWM pulse length |  | RPM* |
| --- | --- | --- | --- |
|  | Forelimb | Hind limb |  |
| SP1 | 1475 | 1525 | 13 |
| SP2 | 1450 | 1550 | 19 |
| SP3 | 1425 | 1575 | 35 |
| SP4 | 1400 | 1600 | 44 |
| SP5 | 1375 | 1625 | 45 |
| SP6 | 1350 | 1650 | 46 |
| SP7 | 1325 | 1675 | 47 |
| SP8 | 1300 | 1700 | 48 |

The final constructions of both the prototype robot (using 3D printed parts from 3D scans) and the necrobot Pak Biawak (using the actual varanid skeleton) are shown in Figure 3. While the gait construction was the same in each, there were minor differences due to post-drying anomalies arising from the varanid skeleton. These included the need to rearticulate the skeleton using a combination of internal metal wiring in combination with mouldable PolyMorph, which meant that all skeletal elements were not in exactly the same position for the necrobot as they were for the 3D printed prototype. The fixings, cf. Figure 1, were designed to allow for a level of variability without compromise in the structural stability of the bots. The electronics housing was additionally moved to a position behind the necrobot, rather than on the pelvic region as in the prototype. This was due to rearticulation with wiring in this region generating higher levels of flexibility in the spine, which was naturally absent in a 3D printed spine.

**Figure 3.:**
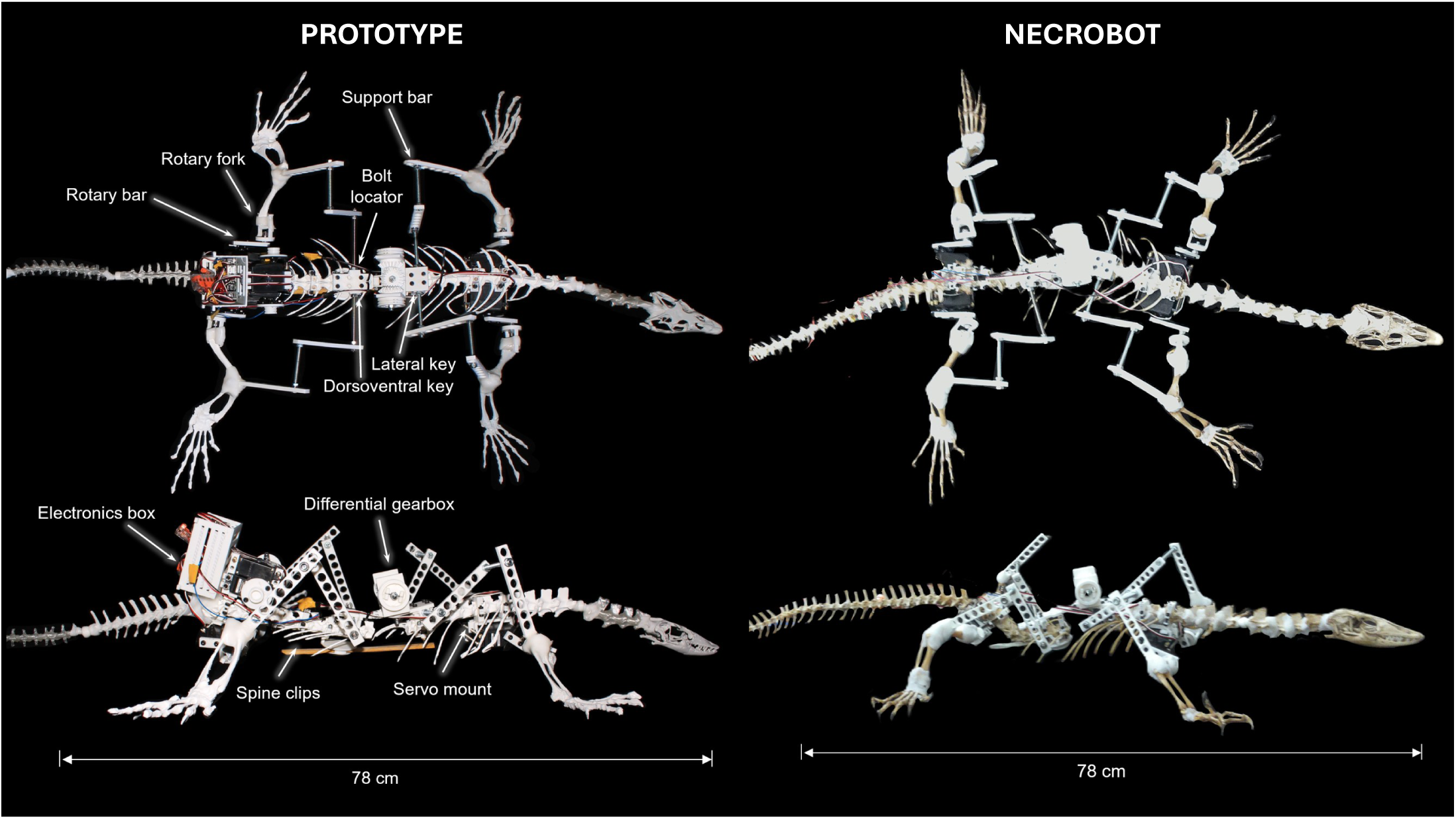
The final constructions of both the prototype robot (using 3D printed parts from 3D scans) and of the necrobot Pak Biawak (using the actual varanid skeleton). The prototype robot was used to enable design of the connectors and fixings for Pak Biawak without damaging the necrobot skeleton.

### 2.2. Trajectory Mapping

Data was extracted using the Tracker: Video Analysis and Modeling Tool [67]. The data extraction process was identical for both the varanid and the necrobot. A total of 62 videos were analysed, 48 for Pak Biawak (the necrobot), and 14 for the live varanid. To extract lateral view data, a reference length was specified at 50 mm for the varanid and 90 mm (representing the length of a bar) for each of the robots. Reference points were chosen for orthogonality to ensure measurement precision. The elbow and knee joints were the two tracked points for the lateral views, shown in Figure 4(a-c).

**Figure 4.:**
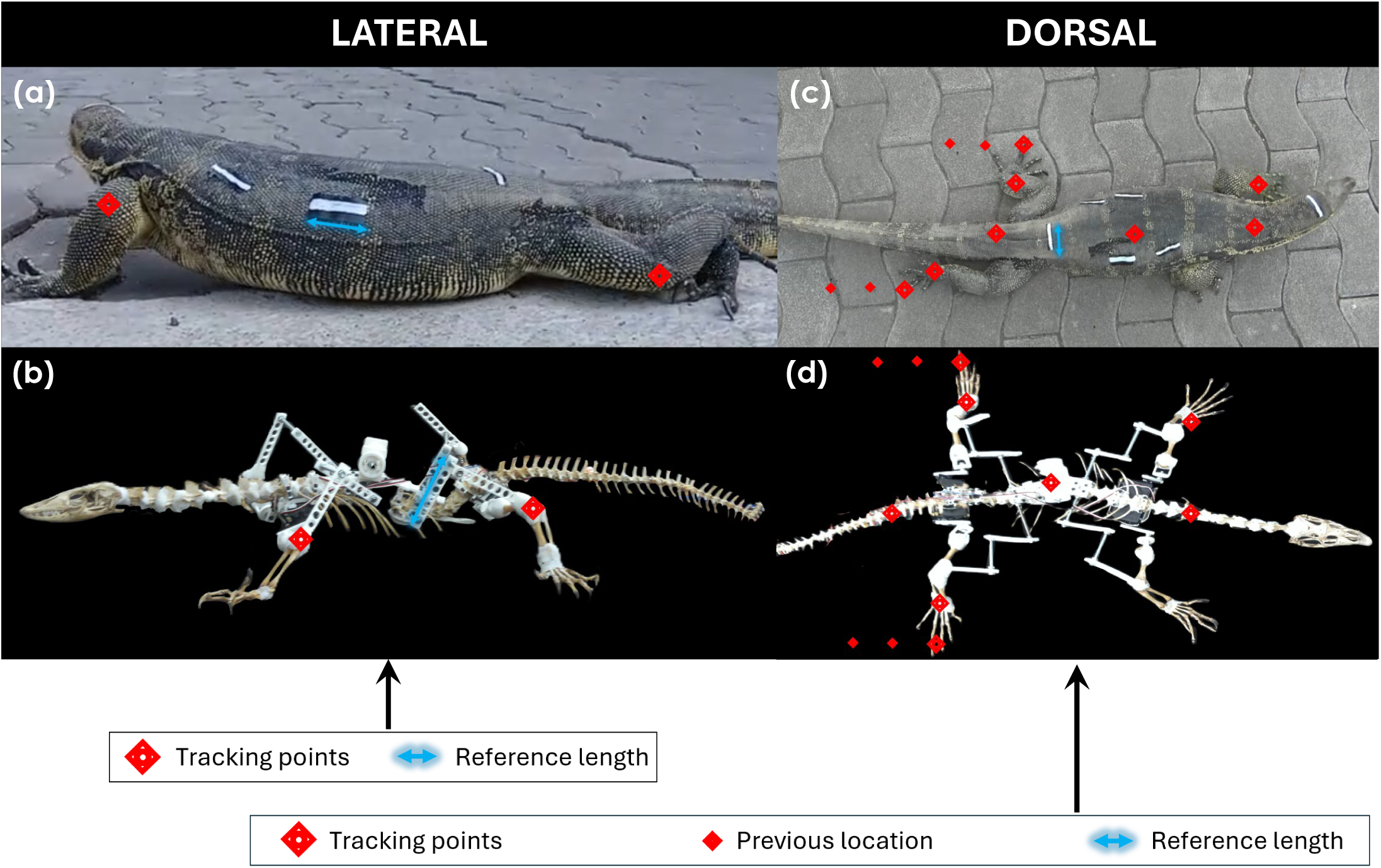
Tracking points in lateral view, with reference length for (a) varanid and (b) necrobot, and in dorsal view, with reference length for (c) varanid and (d) necrobot.

In the dorsal view, reference lengths were specified using 50 mm (using reference length markers) for the varanid and 52.5 mm (representing the skull width) for Pak Biawak. Minimal movement in the robots made them highly susceptible to camera shake, while unpredictable varanid movement caused shifts from the camera orientation. Two reference points were therefore tracked to account for rotation and translation during filming. Seven points were tracked on the body of both the live varanid and Pak Biawak, which are indicated in Figure 4(d-f). The forward and reverse sweeps of the fore and hind limbs were tracked, corresponding to the side captured in the lateral view. The neck, differential and tail were tracked to extract spinal movement. The maximum width during movement was extracted by tracking both hind limbs. All results were exported to text files, then converted into comma-separated values files for processing in Python.

Our focus is on quantifying the consistency of Pak Biawak’s kinematics across different speeds and as compared against, *in-vivo* varanids, which move at varying speeds during locomotion. As such, we will consider the speed-related differences from both lateral and dorsal perspectives. These will then inform our understanding of Pak Biawak’s ability to replicate a live varanid’s sprawling gait.

### 2.3. Lateral kinematics

The elbow and knee joints were tracked in the Cartesian plane to map the two-dimensional lateral gait as discussed in Section 2.2. The trajectory of the elbow was elliptical for both fore and hind limbs. As such, a least-squares ellipse was fitted to the data based on the algorithm outlined in [68], which describes the ellipse-fitting as a minimisation of Equation 1. In this equation, **D** is the decomposed design matrix, **C** the decomposed constraint matrix and the superscript *T* is transpose. Component **a** represents a vector of the coefficients of the general ellipse equation in Equation 2, and **a** is represented by Equation 3. Here, *a, b, c, d, e, f* are coefficients. The least squares minimisation of Equation (1) is achieved by applying Lagrange multipliers, yielding an optimal solution for **a** as shown in the gathered Equations in 4, where **S** is the scatter matrix and *λ* is an eigenvalue.

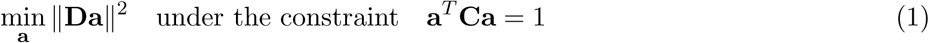

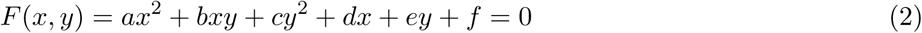

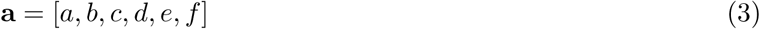

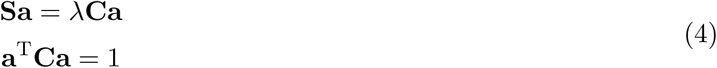

The analysis in [68] yields the equations shown in the gathered Equations in 5, where **M** is the reduced scatter matrix. **C_1_** is the split constraint matrix, **C**; **S_3_** and **S^T^**are components of the split scatter matrix, **S**; and **a_1_** and **a_2_** are the split vector coefficients of **a**. The equations in (5) are equivalent to (4) so can be used to find a suitable eigenvector **a_1_** for **M**.

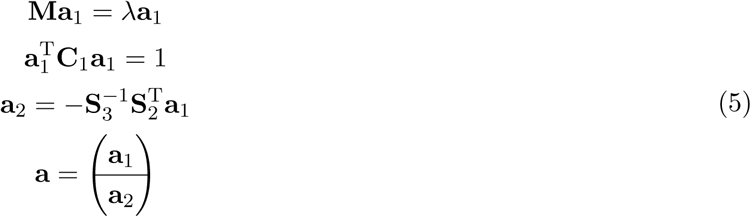

Example plots for the necrobot and varanid are shown in Figures 5 (a) and (b), respectively. Further plots are provided as Electronic Supplementary Material.

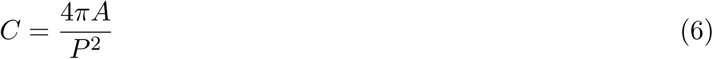

**Figure 5.:**
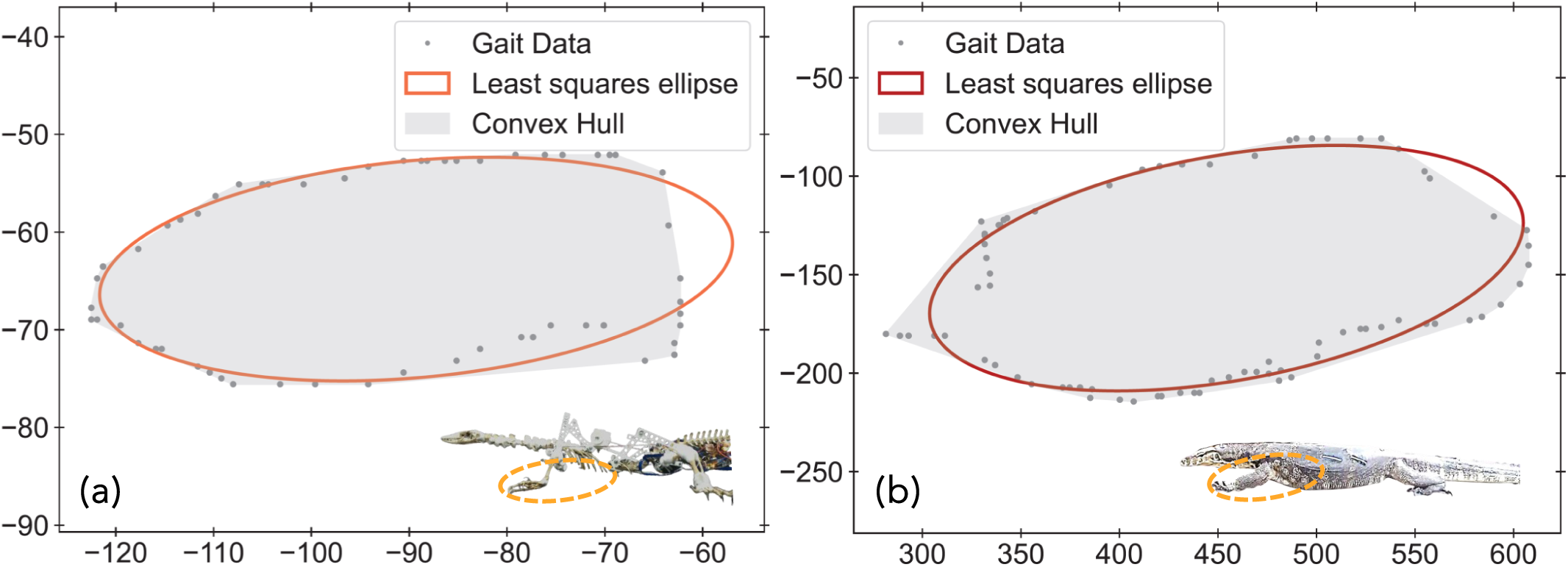
Representative lateral motion plots featuring fitted least-squares ellipse and shaded convex hull for (a) the necrobot (b) the varanid.

We begin by considering ‘shape metrics’ to (a) compare each metric against Pak Biawak’s speed, and (b) compare Pak Biawak’s metrics at all speeds against those of the live varanid. The **aspect ratio** of the fitted ellipse was calculated by dividing the semi-major axis, *r_maj_*, by the semi-minor axis, *r_min_*. The convex hull for the data points was calculated using SciPy’s convex hull library. This finds the smallest convex that encloses all data points, therefore representing the total area covered in one movement. The **normalised area** was calculated by dividing the area of the convex hull by the area of the circle defined by the Feret diameter of the data points. A diagram to illustrate this is provided as Electronic Supplementary Material. Similarly, the **normalised perimeter** was calculated by dividing the convex hull perimeter by the perimeter of the same Feret diameter circle. Normalising both the area and the perimeter from limb movement allows for comparison across datasets with different scales, which is needed here since the live varanid and Pak Biawak do not have the same dimensions. Finally, **circularity**, *C*, of the convex hull was calculated in accordance with Equation 6, where *A* and *P* are the convex hull area and perimeter, respectively. The scatter of data points from the fitted ellipse was determined by calculating the difference between the actual point and the predicted point. A diagram to illustrate this is provided as Electronic Supplementary Material.

The shape metrics are shown in Figure 6 for both fore and hind limbs combined plotting (a) aspect ratio (b) circularity (c) the normalised area and (d) the normalised perimeter. Column (i) in this figure compares the effects of necrobot limb speed only on the shape metric, while column (ii) compares these in (i) against the shape metrics for the live varanid. The mean aspect ratios of Pak Biawak within the speed ranges tested are between ∼0.5 and 2, Figure 6(a-i). The mean aspect ratio of the live varanid lies between these two bounds at ∼1.4, Figure 6(a-ii), which is similar to that of Basilisk lizards (∼1.42) [69] and lacertid lizards (∼2) [70] but is lower than that of the zebra-tailed lizard, which has an aspect ratio of ∼2.96 [71] and that of Bibron’s thick-toed gecko, which (for inclined motion) has an aspect ratio of ∼2 (fore limb) and ∼2.3 (hind limb) [72]. The work of Birn-Jeffery and Higham [72] thus indicates a larger stride in the hind limb, which we find to be also true in Pak Biawak (cf. Electronic Supplementary Material), but not in the live varanid (*V. salvator*). Pak Biawak showed a range of mean aspect ratios between ∼0.4-1.9 (forelimb) and ∼0.4-2.3 (hind limb), while in the varanid, the mean aspect ratio was measured at ∼1.4 in the fore limb and ∼1.3 in the hind limb. The hind limb distribution was nevertheless larger for than the fore limb in the varanid, which indicates that there is greater variability in stride in the hind limb of *V. salvator* (cf. Electronic Supplementary Material). The mean circularity in Pak Biawak considering the full set of speeds, Figure 6(b-i), lies between ∼0.58 and 0.81, indicating the level decircumscribed, and aligning with the measured aspect ratios being *>* 1. The mean circularity of the live varanid is slightly lower than that of Pak Biawak at all speeds (∼0.57), Figure 6(b-ii), however, the distribution is considerably broader for the varanid as it varies gait more freely than Pak Biawak, a characteristic influenced by factors such as energetic and metabolic considerations, microhabitat type, foraging mode, habitat openness [73], and fight/flight behaviour [74]. The swept area from the lateral stroke of a limb in motion has been an important metric to characterise in both robotics [75] and biology [76]. The swept area and its perimeter are defined here as the area and perimeter of the convex hull, respectively, and the normalised area and perimeter calculated by dividing against the area of a circle defined by the Feret diameter of the data points. Figures 6 (c-i) and (d-i), respectively, show these shape metrics at each of Pak Biawak’s speeds. The mean normalised area within the full range of speeds tested is between ∼0.29 and 0.55, while the mean normalised perimeter is between ∼0.72 and 0.82. The equivalent mean normalised area and perimeter for the live varanid are ∼0.25 and ∼0.71, Figures 6 (c-i) and (d-i), respectively, and are thus very close to the range of mean values for this metric exhibited by Pak Biawak.

**Figure 6.:**
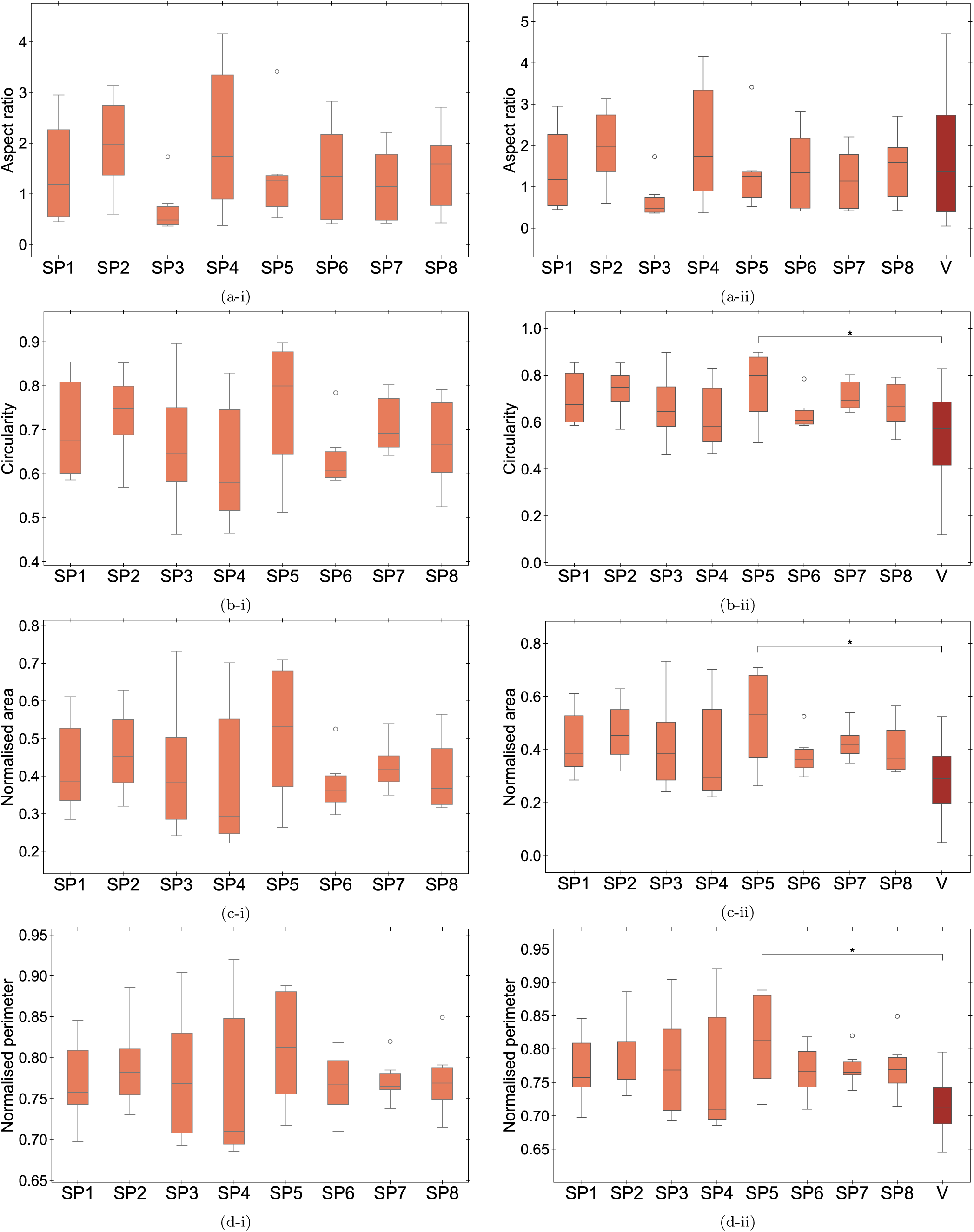
Box and whisker plots of shape metrics across all speeds (SP1-8) for Pak Biawak compared to the varanid (V) for (a) aspect ratio (b) circularity (c) normalised area (d) normalised perimeter, for all limbs. (i) shows the speeds of Pak Biawak, each compared using one way ANOVA and (ii) shows the speeds compared directly against the varanid using Dunnett’s test. Boxes indicate interquartile range (IQR); whiskers extend to 1.5*×*IQR. Significance levels: * p*≤*0.05, ** p*≤*0.01, *** p*≤*0.001; absence of asterisks indicates no significant difference (NS).

When comparing between Pak Biawak’s speeds only (i), we applied a one-way analysis of variance (ANOVA). Pak Biawak’s walking was tested at the eight different speeds (SP1-8), cf. Table 1. The ANOVA tests the null hypothesis, *H*_0_, that the mean, *µ*, of *k* independent groups is equal, Equation 7.

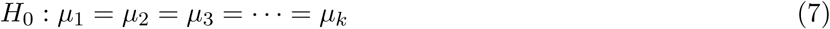

If the analysis returns a p-value *<* 0.05, then the alternative hypothesis is accepted, *H_A_*, which states there is a statistically significant difference between the group means. When comparing the shape metrics at each speed of Pak Biawak directly against the live varanid as seen in column (ii), we apply Dunnett’s tests using the varanid metric as the control group using *α* = 0.05. These tests evaluate whether the mean value of a metric for a specific speed was significantly different to the same metric for the varanid. In both tests, a single asterisk (*) indicates statistical significance at the 0.05 level (0.01 *< p* ≤ 0.05), a double asterisk (**) at the 0.01 level (0.001 *< p* ≤ 0.01), and a triple asterisk (***) at the 0.001 level (*p* ≤ 0.001). While Figures 6(a-d) show the metrics for both fore limbs and hind limbs combined, data and plots for fore limbs only, and hind limbs only, are provided as Electronic Supplementary Material.

When observing the results in column (i) of Figure 6, we find that there are no significant differences (p ≥ 0.05) in any of the shape metrics when comparisons are made across the eight different speeds tested. This indicates that the formed ‘shapes’, as associated with lateral limb movement, are not influenced by the speed at which the limb turns during ambulation. When these are then compared against the same shape metrics for the live varanid column (ii) of the same figure, we note that there are very few examples of difference between the shape metrics of Pak Biawak and the live varanid, with each of these differences arising at one speed (S5) in the circularity, normalised area and normalised perimeter metrics, where p ≤ 0.05.

Our attention now turns to mapping and analysing the ‘trajectory metrics’, which essentially inform us of how closely the kinematic data extracted from the videos follows a predicted path. The general equation of a rotated ellipse is calculated in accordance with Equation 8, while the actual radial distance, *r_a_*, from the centre of the ellipse was calculated using Equation 9, where (*x_c_, y_c_*) is the centre of the ellipse and (*x_a_, y_a_*) are the actual coordinates of a data point.

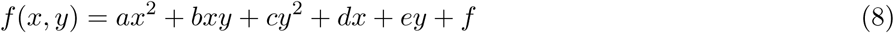

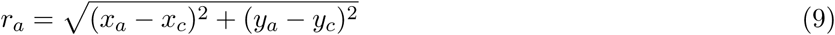

The angle of the actual data point relative to the global horizontal is then computed according to Equation 10.

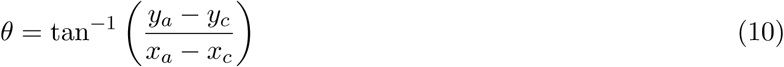

The points on the fitted ellipse can be thought of as the predicted values, denoted by the subscript *p*, with their positions described in polar coordinates. These points lie at the same angle as the actual data point but at some predicted radius, *r_p_*, Equation 11.

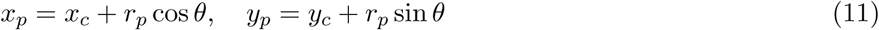

Substituting Equation 11 into Equation 8 yields a quadratic equation for *r_p_*, shown by Equation 12. Here, *a* = *A* cos^2^ *θ* + *B* cos *θ* sin *θ* + *C* sin^2^ *θ*, *b* = 2*Ax_c_* cos *θ* + *B*(*x_c_* sin *θ* + *y_c_* cos *θ*) + 2*Cy_c_* sin *θ* + *D* cos *θ* + *E* sin *θ*, and *c* = *Ax*^2^ + *Bx y* + *Cy*^2^ + *Dx* + *Ey* + *F*; while *A* = ^cos^ ^(*ϕ*)^ + ^sin^ ^(*ϕ*)^, *B* = 2 − cos(*ϕ*) sin(*ϕ*), *C* = ^sin2 (*ϕ*)^ + ^cos2 (*ϕ*)^, *D* = −2*Ax* − *By*, *E* = −*Bx* − 2*Cy*, and *F* = *Ax* + *Bx y* + *Cy*^2^ − 1, where *ϕ* is the rotation of the ellipse.

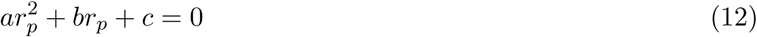

Applying the quadratic formula to Equation 12 yields Equation 13.

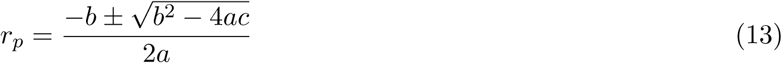

Therefore, for real solutions where *b*^2^ − 4*ac* ≥ 0, the value closest to the actual point can be selected as the predicted radial distance, *r_p_*. The actual and predicted distances were compared using Equation 14.

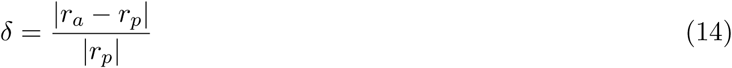

Equation 14 was applied to all *n* points in a dataset, and from this we calculated the **distance mean** (the average radial distance of the ellipse), for each sample set.

The root mean square error (*RMSE*) of the data points from the fitted ellipse was calculated. Considering the fitted ellipse could be rotated by some angle, *ϕ*, points on the ellipse can be described parametrically by Equation 15, where *ψ* is an angle *ψ* ∈ [0, 2*π*] that defines the points on the ellipse.

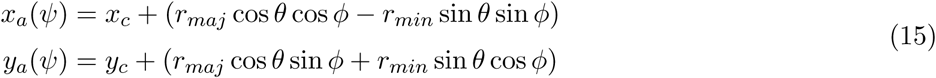

For each data point, the smallest distance between the data point and the ellipse was found by minimising the squared Euclidean distance for each angle *ψ*, Equation 16.

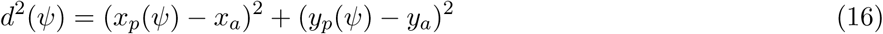

After finding the minimum distance for each point, the *RMSE* was calculated using Equation 17, where *d_i_*is the minimum distance from point *i* to the fitted ellipse.

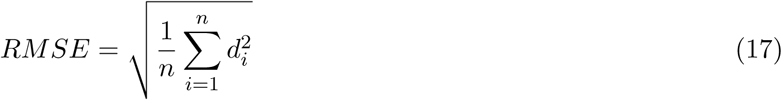

The *RMSE* values were normalised by dividing by the mean distance of the data points from the ellipse centre, *d_mean_*, Equation 18, yielding the **normalised RSME** trajectory metric, *RMSE_normalised_*. Normalising the RMSE allows for comparison across datasets with different scales, which is needed here since the live varanid and Pak Biawak do not have the same dimensions. The trajectory metrics (distance mean, normalised RSME) provide information on the closeness of the actual data to the predicted path.

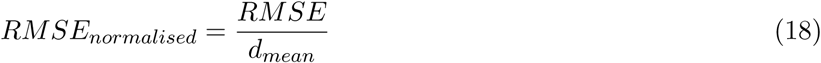

When observing the results in Figure 7 column (i) we find that there are no significant differences in (a) the distance mean and (b) the normalised RMSE metrics, indicating that the trajectory of limb movement is not influenced by the speed at which it turns during ambulation. Further, there is only one instance of a significantly notable difference (p ≤ 0.01) between one of the speeds in Pak Biawak and the live varanid, which occurs at SP3 in the normalised RSME metric. The closeness in the normalised RSME values is important as it implies that there is kinematic consistency in the trajectories of Pak Biapwak at different speeds, and when compared to the live varanid regardless of the natural variance in slip that that either the varanid or necrobot will inevitably experience [77–79]. Kinematic inconsistencies could arise nevertheless, from more extreme cases of slip [80], which could potentially be affected by factors such as body mass and size [81,82].

**Figure 7.:**
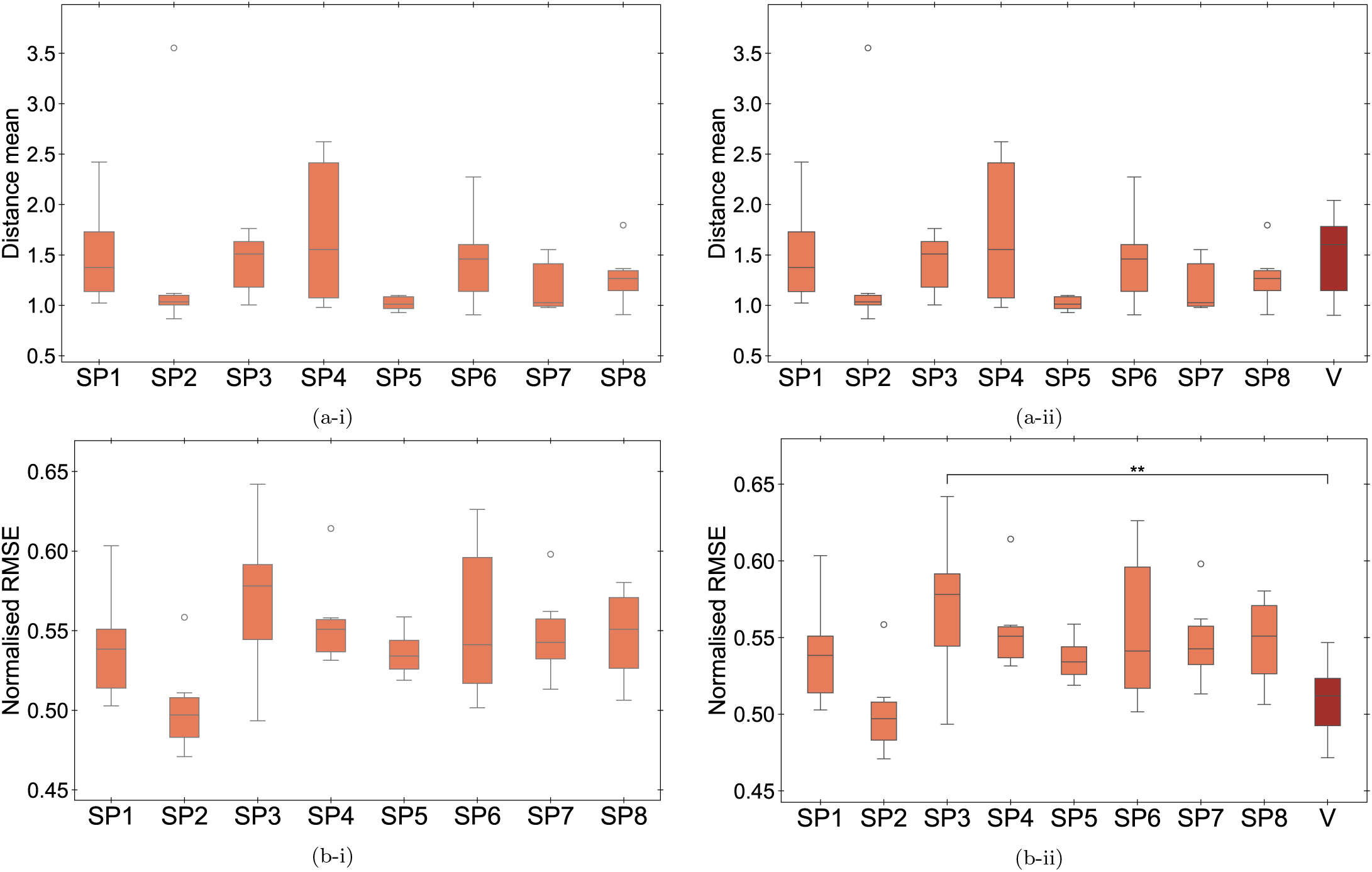
Box and whisker plots of trajectory metrics across speeds (SP1-8) for Pak Biawak only, column (i), and as compared to the varanid (V), column (ii) specifically: (a) the distance mean (average radial distance of the ellipse) and (b) the normalised RMSE, for all limbs. Boxes indicate interquartile range (IQR); whiskers extend to 1.5*×*IQR. Significance levels: * p*≤*0.05, ** p*≤*0.01, *** p*≤*0.001; absence of asterisks indicates no significant difference (NS).

Over the full set of lateral metrics tested (both shape and trajectory metrics), 91.7% of Pak Biawak’s results at all speeds were recorded as having no significant differences over the same metrics of the live varanid. This confirms there is a high level of similarity between the lateral gait of the necrobot Pak Biawak and the lateral gait of the live lizard.

### 2.4. Dorsal kinematics

The changes in spine bending angle during ambulation were calculated by computing the vector between the differential and tail, **v_1_**, and the vector between the neck and differential, **v_2_**, at each time step, Equation 19. The angle between the vectors, *α*, was then calculated using Equation 20.

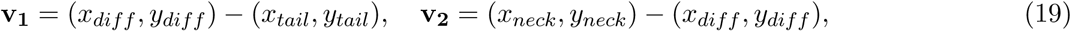

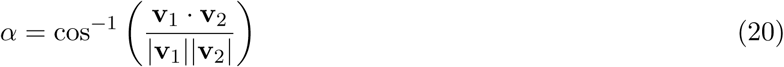

The bending angle, *α*, was normalised by subtracting from the initial angle in the first frame of each video. The spine bending angle was passed through a smoothing Savitsky-Golay filter, using the SciPy ‘savgol filter’. With each data point, the filter considers the points in its neighbourhood window, *N*, and fits a polynomial of specified degree, *M*, using the least-squares method. The key equation, Equation 21, yields the value of the fitted polynomial at the centre point of the current window for the *i*th data point, where *k* is the polynomial power and *a_k_* is the coefficient that is determined when minimising Equation 22. Here, *j* is the relative position of a point to *i* and the total number of points is *M* = 2*n* + 1. A neighbourhood window *N* = 31 and *M* = 3 was selected for calculations.

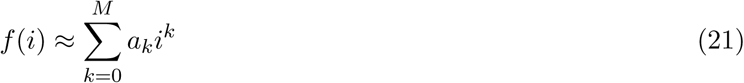

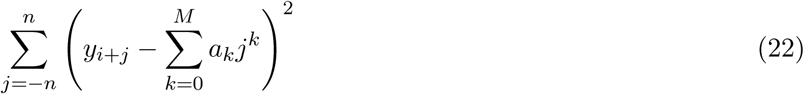

The peaks of the smoothed polynomial were located using SciPy’s ‘find peaks’ function. This finds local maxima by comparing neighbourhood values under specified conditions for the peaks. The prominence of peaks was set to be greater than 0.5 and the width to be greater than 10. The function was applied again to the inverted polynomial to locate troughs.

The moving average of the smoothed polynomial was calculated by finding the x and y midpoints of extrema. These midpoints were then interpolated between to created a plotted moving average. The polynomial **amplitude** in time was calculated by subtracting the extrema from the moving average value. The polynomial **period** was calculated by determining the distance between intersections of the moving average and polynomial. Example plots of the **spine bending angle** for the necrobot and varanid are shown in Figure 8(a) for Pak Biawak and (b) for the live varanid. All plots are provided as Electronic Supplementary Material. Observing the example plots in this figure, we note that the moving average for Pak Biawak delineates from its midline more so than for the varanid and this is indubitably due to the fact that the varnid’s gait is cognitive [83] and decisive, whereas Pak Biawak’s gait is entirely dependent on the geometry of the varanid skeleton. Lizards, like many animals exhibit phenotypic asymmetry [84], and this can be reflected by their gait [85]. In this figure, we note from the amplitude output, that there is more symmetry in the gait of the varanid on its left than on its right, where there is obvious greater variability from the moving average line, which approximates the sagittal plane of the animal during movement. The varanid spine bending recorded here is typical in undulatory form to those of other sprawling lizards [86], and is a result of differential leg movement during sprawling.

**Figure 8.:**
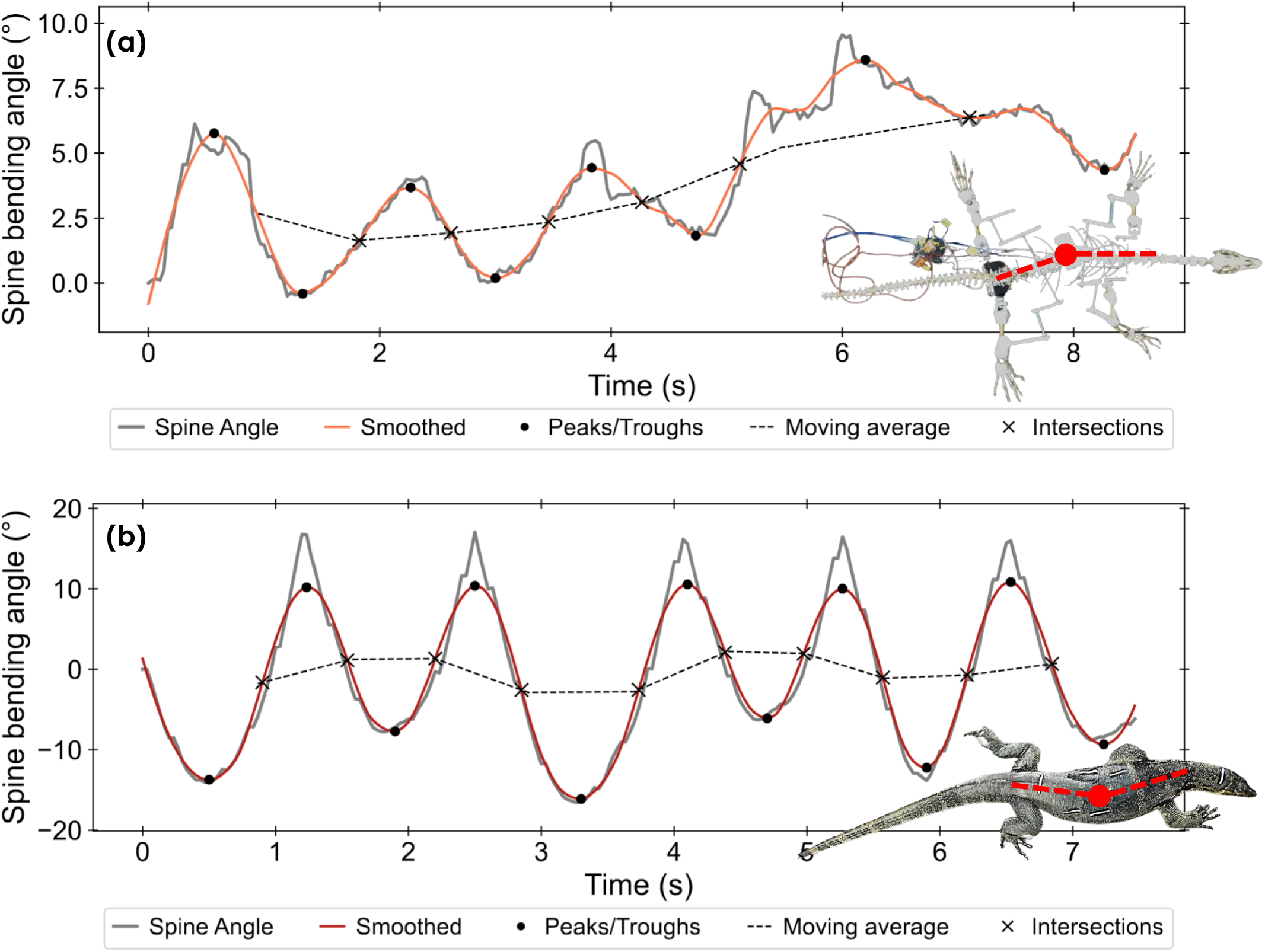
Example plots of spine bending angle vs time including Savitsky-Golay filtered polynomial, indicated extrema, moving average line and indicated intersections for (a) necrobot and (b) varanid.

Figures 9(a-i) and (b-i) show the box and whisker plots for Pak Biawak’s mean amplitude and average period, respectively. Significance bars (one way ANOVA testing) are provided in the plots at the following significance levels: * p ≤ 0.05, ** p ≤ 0.01, *** p ≤ 0.001. The absence of significance bars and asterisks indicates there is no significant difference between the data sets. Here, we see that while the average periods are not statistically different for all speeds (SP1-8), the mean amplitude shows there are significant differences (p ≤ 0.05) primarily between SP1 and SP3, SP4 and SP5, with an additional pairwise difference (p ≤ 0.05) observed between SP3 and SP8. The high variability of the amplitude between speeds reflects the higher level of complexity involved in designing a sprawling necrobot with repeat passive spine bending movements, since spine bending is itself affected by the start and finish positions of the spine during ambulation [87], and there are no active mechanisms [88] to restore posture at key stages of limb rotation. When each of the metrics in (a-i) and (b-i) at the different speeds are compared (using Dunnett’s test) against the mean amplitude and mean period of the live varanid, Figures 9(a-ii) and (b-ii), respectively, we confirm there are no statistically significant differences that can be highlighted between any of Pak Biawak’s metrics at any speed, and the equivalent live varanid metric.

**Figure 9.:**
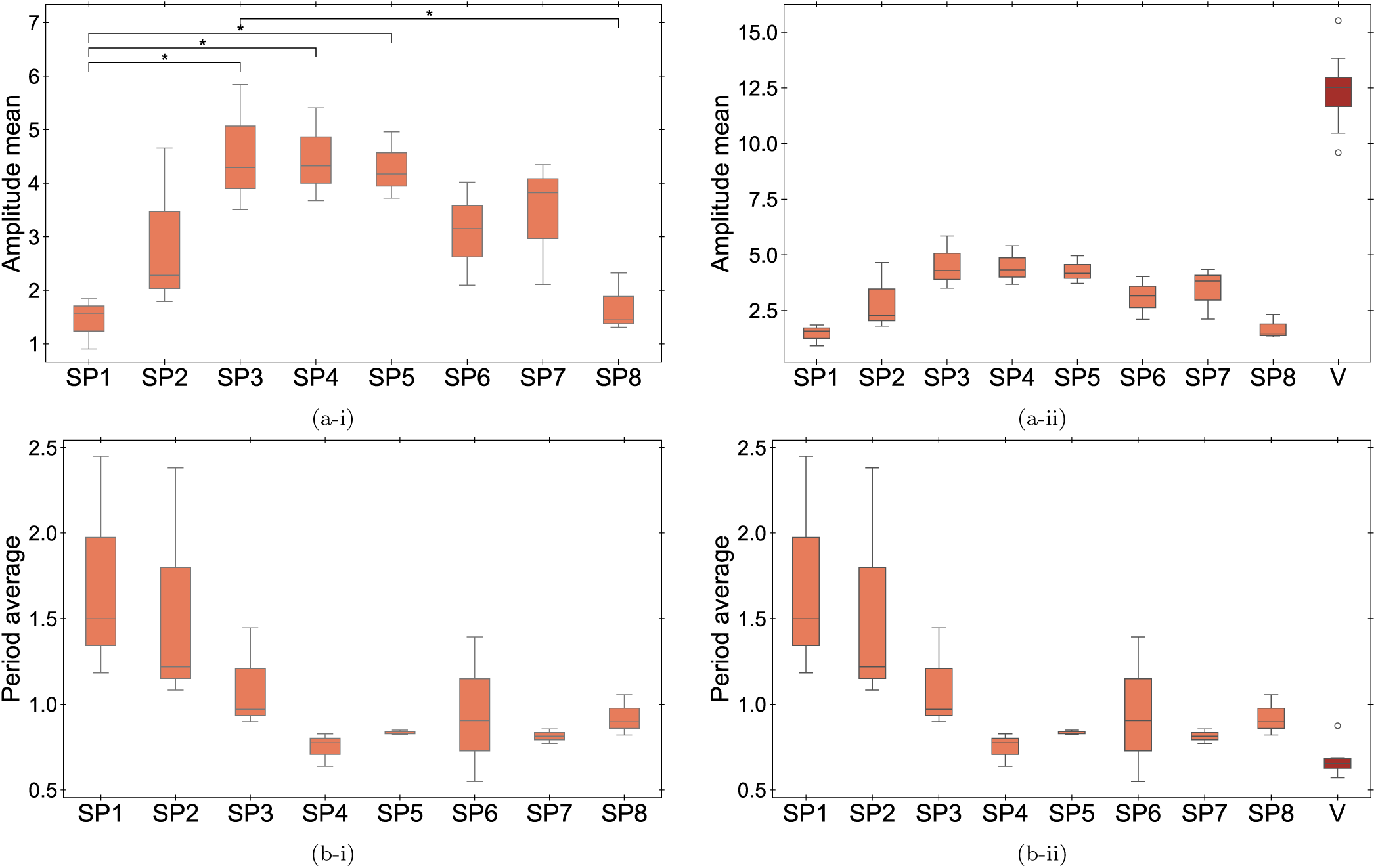
Box and whisker plots comparing (a) mean amplitude and (b) average period across speeds (SP1-8) of the necrobot against the varanid (V), (i) compares the speeds only and (ii) compares the metrics at each speed against those of the live varanid. Boxes indicate interquartile range (IQR); whiskers extend to 1.5*×*IQR. Significance levels: * p*≤*0.05, ** p*≤*0.01, *** p*≤*0.001; absence of asterisks indicates no significant difference (NS).

We redirect our attention to limb motion, albeit from a dorsal perspective. Representative plots of the forward and reverse sweeps from Pak Biawak and of the live varanid are shown in Figure 10, revealing generic sweep trajectories [9] during ambulation. All plots are provided as Electronic Supplementary Material. A second order polynomial was fit to the data of forward and reverse sweeps for limbs using NumPy’s ‘polyfit’ function. This function seeks to minimise the least-squares error, *E*, Equation 23, where *p*(*x_j_*) is the polynomial predicted value at position *x_j_*, *y_j_* is the actual data value at position *x_j_* and *k* is the number of data points minus one.

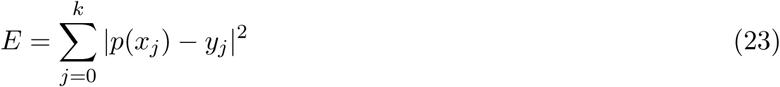

**Figure 10.:**
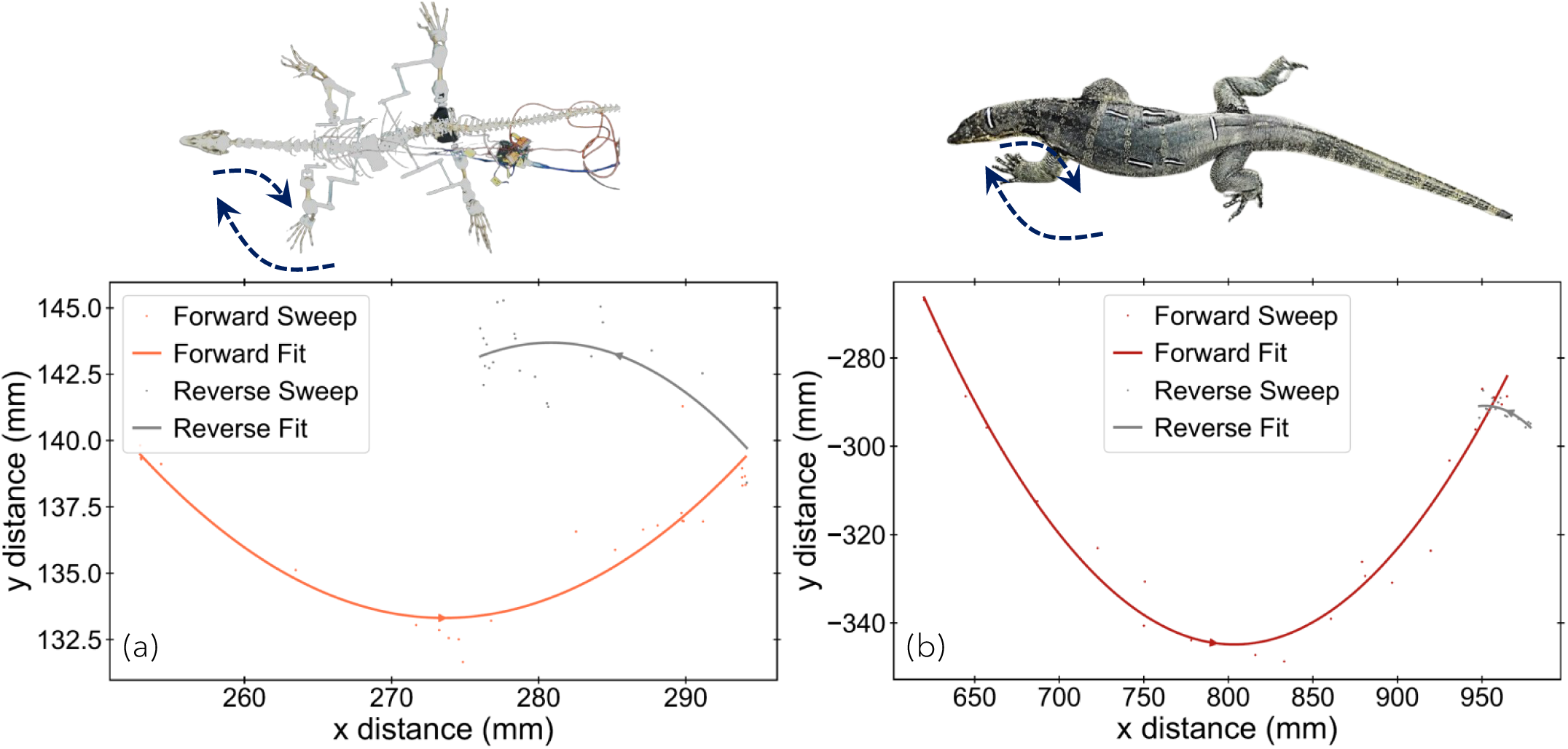
Example plots of x-distance vs. y-distance moved during limb sweep for (a) necrobot forelimb and (b) varanid forelimb.

The **angular curvature** of the fitted polynomial to the forward and reverse sweeps was calculated. Similar to Equations in 19, two vectors were calculated using the polynomial start, middle and end points. The angle between the vectors was computed using Equation 20. An illustration depicting this is provided as Electronic Supplementary Material. The differential curvature, *κ*, or, **curvature**, at the midpoint of the fitted polynomial was calculated using Equation 24, where *f^′^*(*x*) and *f^′′^*(*x*) are the first and second derivatives of the fitted polynomial. This gives the reciprocal of the radius of the osculating circle at the midpoint, *R*. An illustration is provided to depict this as Electronic Supplementary Material.

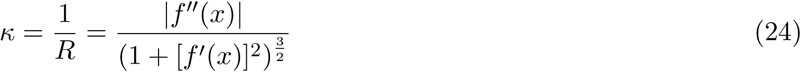

Figures 11(a-i) and (b-i) show Pak Biawak’s angular curvature in forward and reverse, respectively, at different speeds for both fore and hind limbs combined. The mean angles range from ∼120*^◦^* to 175*^◦^* in the forward sweep, and from ∼135*^◦^* to 170*^◦^* in the reverse sweep. Aligned concepts such as pace angulation,_(_⟨*P* ⟩, _)_have been the stride length while *w_t_* is the trackway width. Further, on steep and vertical inclines, climbing affects stride angle as reported by Schultz et al. [90] who considered the kinematics of house geckos. Here, we also note that there are generally no statistically significant differences in either forward or reverse angular curvature during a sweeping limb with the exception of notable differences (p ≤ 0.05) between SP8 against SP5 and SP6 for the forward sweep, Figure 11(a-i).

**Figure 11.:**
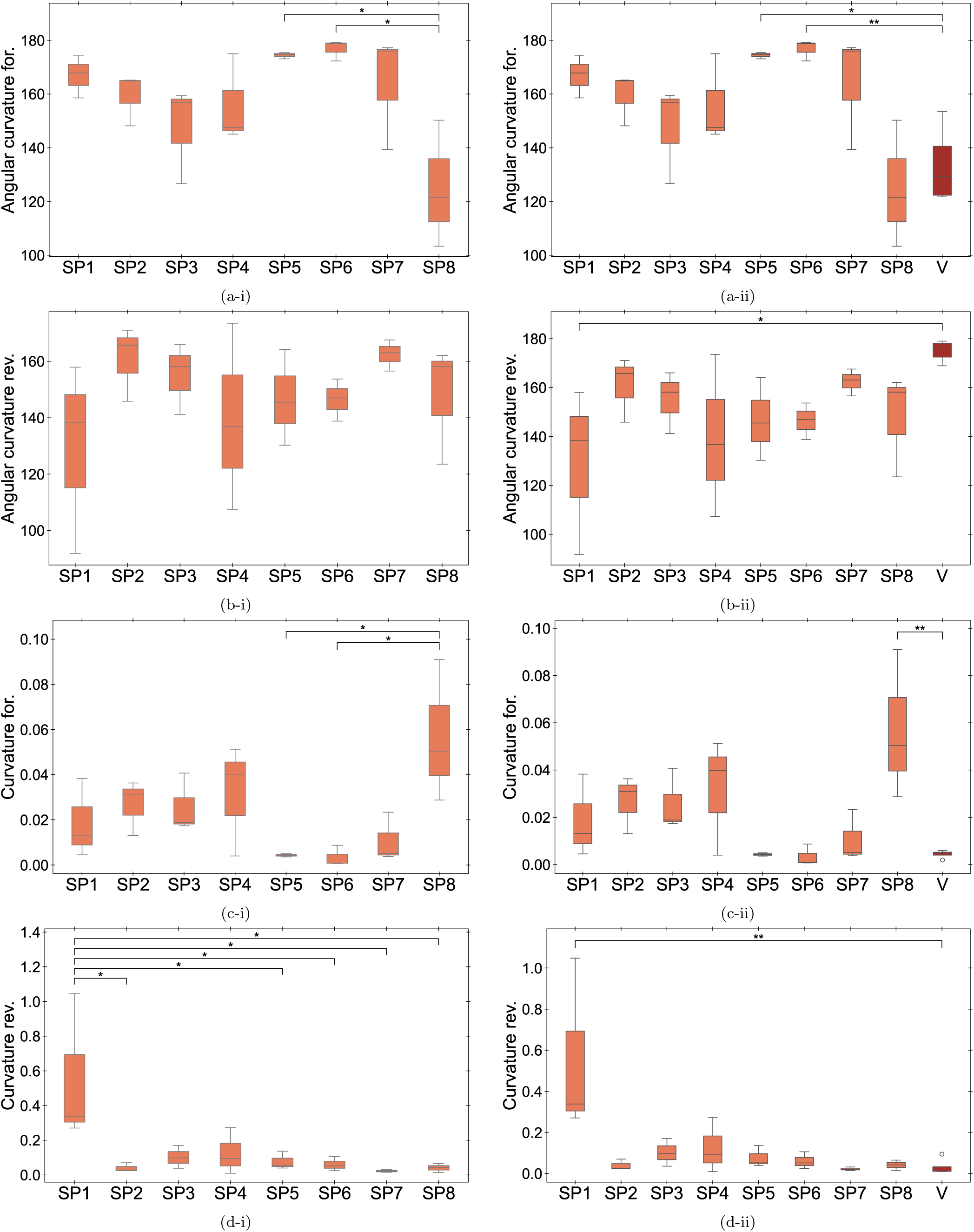
Box and whisker plots of curvature metrics across speeds (SP1-8) compared to varanid (V) for (a) angular forward (b) angular reverse (c) differential forward and (d) differential reverse where (i) hows comparison of speeds only and (ii) shows comparisons against the varanid. Boxes indicate interquartile range (IQR); whiskers extend to 1.5*×*IQR. Significance levels: * p*≤*0.05, ** p*≤*0.01, *** p*≤*0.001; absence of asterisks indicates no significant difference (NS).

When considering the curvatures in forward and reverse, Figures 11(c-i) and (d-i), respectively, we find that the same notable differences (p ≤ 0.05) exist between SP8 against SP5 and SP6 for the forward sweep (c-i), while the slowest speed (SP1) shows the most significant differences (p ≤ 0.05) from the majority of the rest of the group in the reverse sweep (d-i). There is overall nevertheless, a preponderance of similarities in both the angular curvatures and curvatures between most sample sets. The mean curvatures in forward and reverse range from ∼0 to 0.05 and ∼0 to 0.3, respectively. The closer the curvature is to 0, the closer the limb sweep trajectory is to a minimal surface boundary, which we find is overall more evident in Pak Biawak’s forward sweep than in its reverse sweep.

We next use Dunnett’s test to compare both forward and reverse angular curvatures and curvatures at each speed against the equivalent metric of the varanid, which has a mean angular curvature of ∼130*^◦^* and ∼175*^◦^*in forward and reverse sweeps, respectively. Forward and reverse angular curvatures, Figure 9(a-ii) and (b-ii), respectively, reveal considerable similarity between the angular curvatures computed for the live varanid gait with those of Pak Biawak at most speeds. The only instances where significant differences are noted are SP5 (p ≤ 0.05) and SP6 (p ≤ 0.01) against the varanid in the forward sweep, and SP1 (p ≤ 0.05) against the varanid in the reverse sweep. When comparing the Pak Biawaks’ curvatures against those of the live varanid in the same way for forward and reverse sweeps, Figure 9(c-ii) and (d-ii), respectively, we find that there is again, considerable similarity between them with the only exceptions being between SP8 and the live varanid (p ≤ 0.01) in the forward sweep and SP1 and the live varanid (p ≤ 0.01) in the reverse sweep. All data from Figure 11 separated into fore limb and hind limb only, as well as combined, are provided as Electronic Supplementary Material.

The arc length is the final limb sweep trajectory we will consider. This was calculated from the fitted polynomial to the forward and reverse sweeps by numerical integration, Equation 25, where *a* and *b* are the start and end *x* positions of the arc. The integral was solved using SciPy’s ‘integrate.quad’ function. The arc length was normalised (**normalised arc length**) against the shoulder-to-hip length, which was 350mm for Pak Biawak and 320mm for the live varanid. Normalisation of this metric is important for the sake of comparability, as the size differences between Pak Biawak and the live lizard will inevitably affect the arc length of a sweeping limb during ambulation.

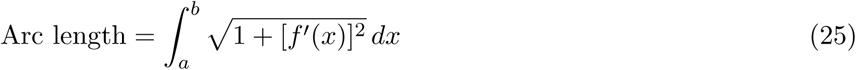

When comparing Pak Biawak’s forward and reverse arc lengths at different speeds using a one way ANOVA test, Figure 12(a-i) and (b-i), respectively, we find that there is no statistical difference in arc length arising as a function of ambulation speed, except in the case of the reverse sweep, where the two highest speeds are each significantly different to the lowest speed (p ≤ 0.05). When using Dunnett’s test to compare Pak Biawak’s forward and reverse arc lengths against that of the live varanid, Figure 12(a-ii) and (b-ii), respectively, we find that Pak Biawak’s arc lengths at every speed are significantly different (p ≤ 0.001) to that of the live varanid for the forward sweep, but that there are no statistical differences at any speed for the reverse sweep. A reason for this could be attributed to combination of lateral spine bending coupled to a lateral limb movement in the live lizard in the forward sweep. This is very different to the forward arc in Pak Biawak, which only has rotational limb motion in the dorso-ventral plane of the necrobot. Pak Biawak is able to generate an arc length only through passive spine movement, which is low. As such, while Pak Biawak is able to exhibit a sprawling gait in its forward sweep, it is unable to mimic the that of the live lizard in terms of its relative arc size. The reverse arc is likely similar in both Pak Biawak and the live lizard because in both, the reverse is very much a push and drag, rather than an extended arc.

**Figure 12.:**
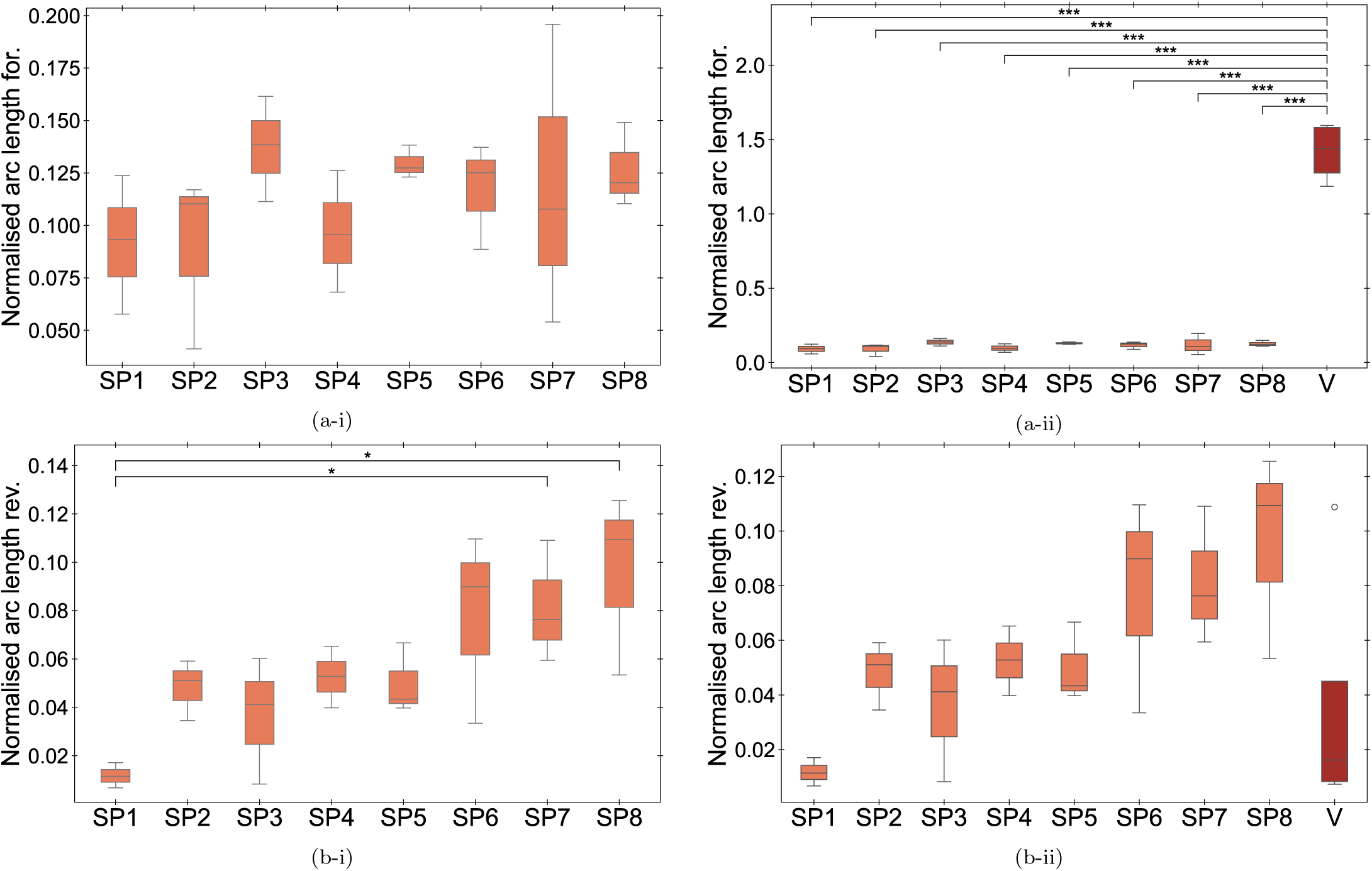
Box and whisker plots of normalised arc length metrics across speeds for Pak Biawak only, column (i) and as compared to the live varanid (V), column (ii): (a) forward sweep and (b) reverse sweep. Boxes indicate interquartile range (IQR); whiskers extend to 1.5*×*IQR. Significance levels: * p*≤*0.05, ** p*≤*0.01, *** p*≤*0.001; absence of asterisks indicates no significant difference (NS).

An important aspect of recording each of the angular curvature (stride angle), the curvature and the normalised arc length, is that using these three metrics provides a multi-tier perspective of dorsal limb sweeps in either lizards or robots in a sprawling gait, building on classical perspectives reported in previous studies that map Cartesian sweep trajectories [70] and which consider stride angles [90]. Here we note that both the angular curvature and the curvature are consistent metrics to use at most speeds in both forward and reverse, aligning also with the those of the live varanid, whereas the arc length, while consistent for the necrobot, only aligns with the live varanid in the reverse sweep and as such, appears harder to mimic. This could be due to limitations to necrobot motion, which is constrained in its kinematic freedom by the simplicity of its construction confining it to single plane leg rotations coupled to passive spine bending at only a singular point on the spine.

Our final gait analysis in the dorsal view relates to the relative trackway width of the sprawling gait for both Pak Biawak and the live varanid. To enable this analysis, individual data points mapping the maximum gait trackway width were connected by linear interpolation. The maximum value was normalised by a nominal width. The nominal widths were 600 mm for Pak Biawak and 430 mm for the varanid and the mean measured across one entire sample in a sample set represents the **normalised difference mean**. Representative plots of the maximum width trace are shown for the necrobot and varanid in Figures 13(a) and (b), respectively. All plots are provided as Electronic Supplementary Material. These width metrics are influenced by all trajectories (limbs and spine) from the sprawling gait, and are shown as box and whisker plots in Figures 13 specifically the normalised difference mean (c) at each of Pak Biawak’s speeds and (d) as a comparison between Pak Biawak’s speeds against those measured for the live varanid. Using a one way ANOVA test in Figure 13(c), respectively we find there are no statistically significant differences in the normalised difference mean at any of Pak Biawak’s speeds, while Dunnett’s tests comparing each of these against the normalised difference mean of the live varanid concurrently show that there is no difference between them, Figure 13(d). These findings indicate there is regularity in the overall relative sprawling gait width for both the necrobot, Pak Biawak, and the live varanid. Sprawling is a specific gait that occurs when limbs protrude laterally from the body and animals with this gait tend to have a wider trackway width than animals that are more erect or that have a parasagittal stance [89,91]. By confirming here that the trackway width of Pak Biawak is similar at all speeds to that of the live varanid, we confirm that the gaits are aligned from the perspective of this final gait metric.

**Figure 13.:**
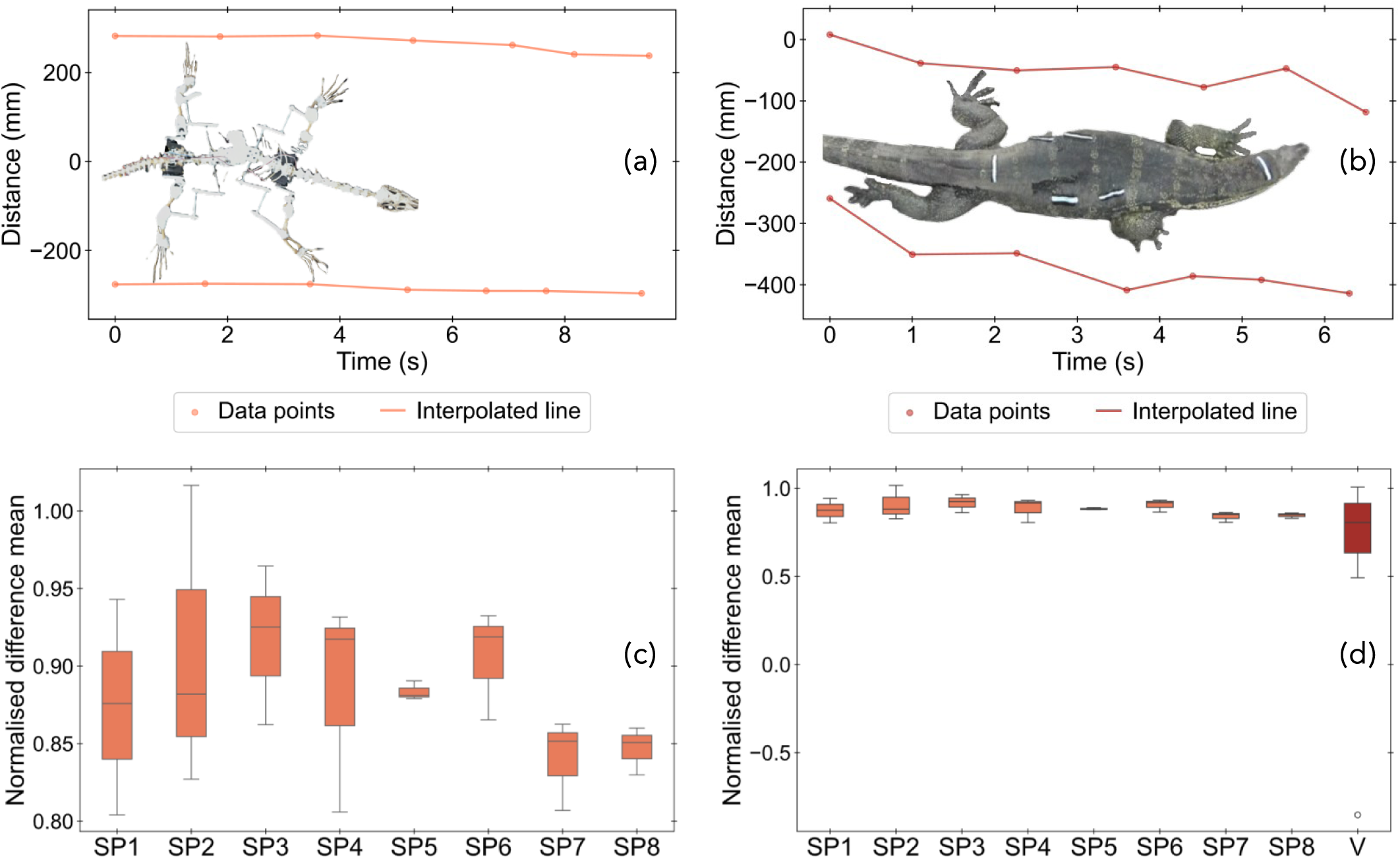
Representative plots of y-distance vs. time tracking the maximum width during walking for the (a) necrobot and (b) varanid. Box and whisker plots of normalised width difference metrics across speeds (SP1-8) for (c) Pak Biawak only and (d) as compared to the live varanid

(V). The absence of asterisks and significance bars indicates there is no significant difference (NS) between any of the sample sets in either (c) or (d).

## 3. Conclusions

The primary goal of this study was to explore the potential for a necrobot fabricated from a varanid (*V. salvator*) skeleton in replicating the complex, sprawling *in-vivo* gait of a varanid from the same species and of approximately the same size. A prototype robot was designed and manufactured to enable the iterative design and testing of adaptable parts for use on the final necrobot. The necrobot, Pak Biawak was assembled by rearticulating the skeleton and attaching the parts designed on the prototype, to the skeletal system of the decreased varanid. Both Pak Biawak (the necrobot) and live varanid were filmed to characterise the kinematics of their sprawling gait. Kinematic metrics were calculated in both lateral and dorsal views, and these were analysed statistically, to examine differences between the two. The necrobot, Pak Biawak, demonstrated strong gait imitation to the live varanid across several different speeds. This is particularly notable when assessing lateral view shape metrics, which include: stride aspect ratio, stride circularity, normalised stride swept area, and normalised stride swept area perimeter. Pak Biawak also has comparable lateral trajectory metrics at different speeds to those of the live varanid including, the radial distance of swept area, and normallised root mean squared error. From a dorsal viewpoint, metrics such as spine bending amplitude and period were found to not be significantly different to those of the live varanid, however, we note that Pak Biawak’s amplitude is affected by sprawling speed. Three metrics were used to compare forward and reverse limb sweeps. These include, angular curvature, differential curvature, and a normalised arc length. From these metrics, several highly significant differences (p ≤ 0.001) were observed when comparing specifically the forward sweep arc length of Pak Biawak at every sprawling speed against the forward sweep arc length of the live lizard. All other kinematic metrics in the necrobot were nevertheless, notably close to those of the live lizard. The trackway width of Pak Biawak was found to be kinematically compatible when compared against the live lizard. Our work demonstrates the feasibility of using necrobot systems with relatively simple actuation to model complex biological locomotion such as sprawling. Our work also potentialises the use of bones as renewable, sustainable and biodegradable structural elements in robotics, providing an alternative to traditional plastic or metal robotic parts.

## 4. Materials and methods

### 4.1. Animal selection

*Varanus salvator*, commonly known as the Asian water monitor, was selected as the varanid species of interest due to its availability in Indonesia, as well as for its size. *V. salvator* lizard is considered to be the second largest lizard species after the Komodo dragon [92]. The large skeleton size of *V. salvator* alongside its relatively slow gait allows for the easy integration of actuating mechanisms into its skeleton to form a necrobot. The animal was sourced and euthanised at the Universitas Gadjah Mada, Yogyakarta, Indonesia following ethical clearance processes.

### 4.2. Skeletal preparation and characterisation

The dead animal was defleshed and its skeleton cleaned by boiling. Some sections of the skeleton remained intact after boiling due to presence of connective tissue. These sections were kept whole to enable easier 3D scanning for a 3D printed prototype, which would be used for the design, testing and placement of connectors, joints and components. An identification system, Figure 14, was developed to help organise the skeleton and any subsequent data. Each skeleton section was placed onto a non-reflective black sheet and photographed using a Canon EOS 1100D. Examples of the original bone images are provided as Electronic Supplementary Material. Key characteristics of skeleton segments were measured using analogue vernier callipers and weighed on an Ascher digital scale. This data is provided in detail as Electronic Supplementary Material. These dimensions were later used to verify the accuracy of 3D printed part, and for calculating the torque requirements of the actuation systems.

**Figure 14.:**
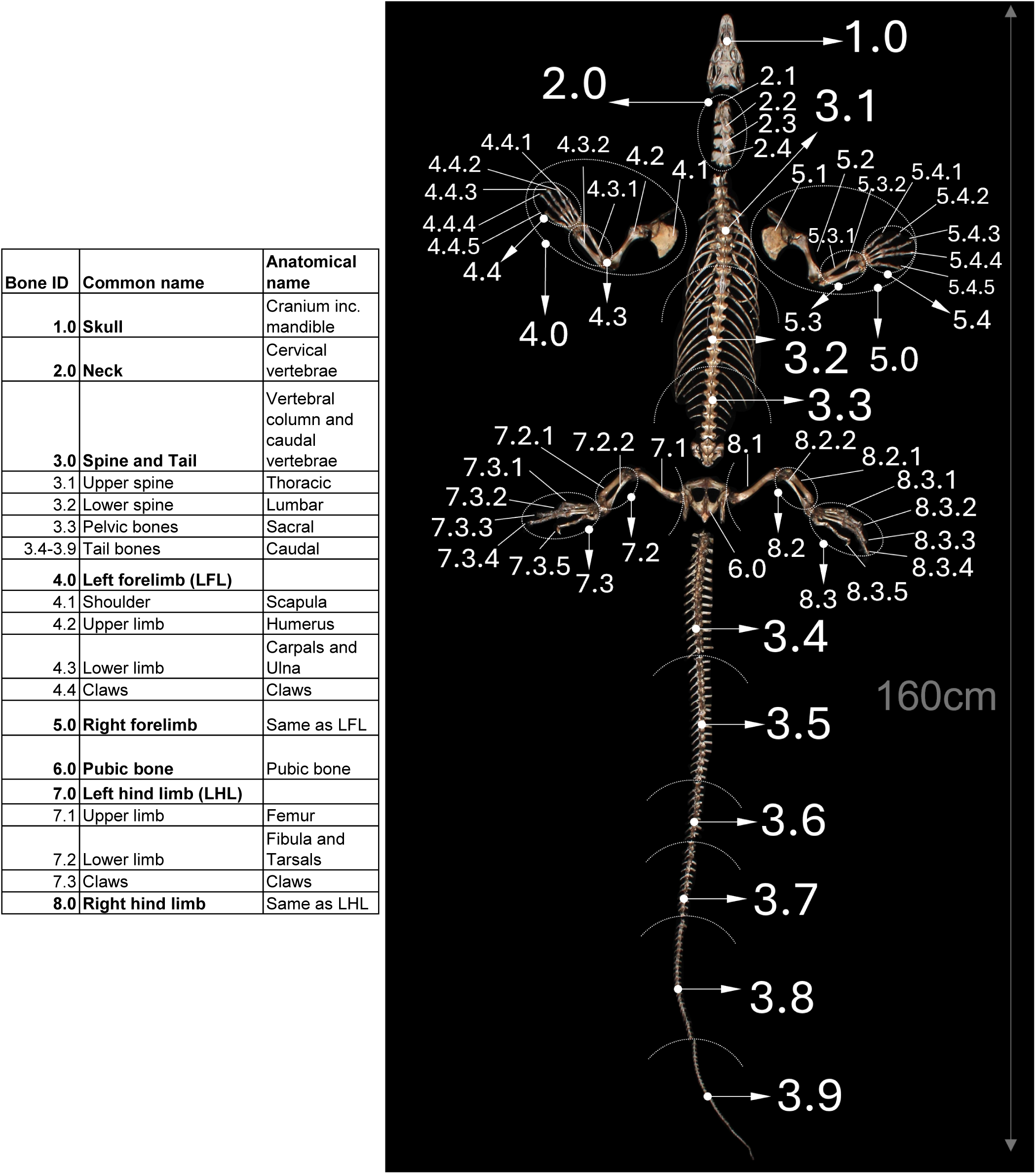
ID of varanid skeletal elements accompanying a labelled diagram of the varanid skeleton.

### 4.3. 3D scanning

3D scanning was conducted with the Revopoint Mini 3D scanner (Revopoint 3D Technologies Inc., Shenzhen, China), which uses structured blue light technology to create point clouds [93]. The scanner was placed on a stable platform at an optimal distance automatically calculated by the Revo Scan 5 software (Revopoint 3D Technologies Inc., Shenzhen, China). Skeletal parts were affixed to a Mcbazel 360 degree rotating turntable (Mcbazel, Lai Chi Kok, KLN, Hong Kong) using putty, and each part scanned over a full rotation. To maximise the scan quality, each scan was conducted using a non-reflective sheet as a backdrop. Larger skeletal parts were scanned in sections, with the part base affixed to the turntable and the top supported using taut black thread. This enabled a steady axial rotation of the part as driven by the turntable, during which a full rotational scan of one section could be completed. For larger parts, multiple complete scans were performed, ensuring significant overlap with previous scans. The overlapped sections were were then used to merge the separate scans together to form a single computational render of a part. In cases where automatic merging was unsuccessful, merging was completed by manually choosing reference points on the skeleton to connect. Meshes were generated using the Revo Scan 5 software and exported as stereolithographic (.stl) format for further post-processing in MeshMixer (Autodesk, San Rafael, California, USA). The Inspector tool was used to automatically detect and correct issues such as disconnected floating objects and mesh holes. Any sections that were not captured accurately, such as the claws, were refined using the rounding tool to improve their shape for 3D printing. To ensure consistency, the surface of the mesh was expanded uniformly in areas matching those where earlier measurements were made, thus minimising inaccuracies in the final computational render.

### 4.4. 3D printing

3D printing by the fused deposition method (FDM) was used to manufacture both the full structure of the prototype, and the kinematic and connector parts for Pak Biawak (the necrobot). The prototype was manufactured to enable design and testing of parts and components prior to their application to the real varanid skeleton. We used the Original Prusa MK4 (Prusa Research, Prague, Czech Republic) and the Kingroon K3PS (Kingroon Tech. Co., Shenzen, China), printing with white PLA filament. The printers had similar resolution so their impact on the structural integrity of the parts was minimal. Large connected sections of the skeleton were separated in judicious locations to enable the integration of actuation mechanisms. The spine for example, was halved to facilitate lateral spine bending through a differential geared connection mechanism. Print orientation was optimised to ensure that the expected loading would occur perpendicular to the printed filament lengths, following Yao et al. [94].

### 4.5. Videography

The varanid and necrobot were individually filmed in both lateral and dorsal views. Due to the unpredictability of the varanid gait and velocity, several repeat videos were taken of the varanid walking. In addition, to extract kinetic and kinematic data using tracking tools, 5 cm strips of electrical tape were placed on the back and sides of the varanid, which were as reference lengths during data extraction. The necrobot was filmed moving at eight different walking speeds, using three repeats per speed, but with multiple complete leg rotations in each video capture. The walking speed was controlled by servo motor rotation over a one second window (cf. Table 1). The robot was filmed in front of a non-reflective backdrop and for lateral filming, the camera was stabilised by placing it on a wheeled platform that moved laterally at the robot walking speed. During dorsal filming, the robot was placed on a non-reflective sheet above the centre of the robot and the camera position maintained relative to the robot while walking.

### Electronic Supplementary Materials

1. Electronic Supplementary Materials (main pdf document contains: (1) Images of skeleton parts (2) Skeleton section masses (3) Lateral trajectory plots for necrobot and varanid (4) Illustration of convex hull normalisation (5) Illustration of scatter distance calculations from the fitted curve (6) Comparison of necrobot lateral metrics over different speeds: all plots (7) Comparison of prototype and necrobot lateral metrics against varanid lateral metrics at different speeds: all plots (8) Angular and differential curvatures (9) Dorsal spine bending all plots (10) Dorsal limb sweep all plots and (11) Dorsal maximum trackway width trace all plots)
2. Electronic Supplementary Video: Pak Biawak sprawling lateral view
3. Electronic Supplementary Video: Live varanid sprawling lateral view
4. Electronic Supplementary Video: Pak Biawak sprawling dorsal view
5. Electronic Supplementary Video: Live varanid sprawling dorsal view

## Supporting information

Electronic Supplementary Materials and Videos

## Data Availability

Data is available from the corresponding author on reasonable request, and has also been included as supplementary material to this paper.

## Ethics Statement

All animal care, handling, and experimental procedure in this study were carried out in out in accordance with The Committee of Ethical Clearance for Pre-clinical Research of The Integrated Laboratory of Research and Testing, Gadjah Mada University, Yogyakarta, Indonesia and an ethical clearance certificate has been granted (Ref. No.: 00014/III/UN1/LPPT/EC/2025).

## Author Contributions

Conceptualization (PA); Data curation (LF, DSY, PA); Formal analysis (LF, PA); Funding acquisition (LF, PA); Investigation (LF, DSY, PA); Methodology (LF, DSY, PA); Project administration (DSY, PA); Resources (DSY, PA); Software (LF); Supervision (PA); Validation (LF, DSY, PA); Visualisation (LF, PA); Roles/Writing - original draft (LF, PA); Writing - review and editing (LF, PA).

## Acknowledgment

We wish to thank the following individuals for their support over the course of this project. FX Sugiyo Pranoto (Pak Frans) affiliation: Museum of Biology, Faculty o Biology, Universitas Gadjah Mada. The following Master of Science students of the Faculty of Biology: Ananto Puradi Nainggolan, Maula Haqul Dafa, and Rashif Naufal Andika. The following Bachelor of Science students of the Faculty of Biology: Afif Fatah Rizki, Arkanniti Dibyawedha Adisajjana, Anthera Al Firdaus Prissandi, Intan Lutfianawati, and Salman Ali Nazar. The monitor lizard caretaker: Ari Irawan.

