## Supplementary material for "Gait analysis of Pak Biawak: a necrobot lizard built using the skeleton of an Asian water monitor (*Varanus salvator*)": Electronic Supplementary Materials and Videos: Electronic Supplementary Material.pdf

**ARTICLE HISTORY**

Compiled July 11, 2025

### 1. Images of skeleton parts

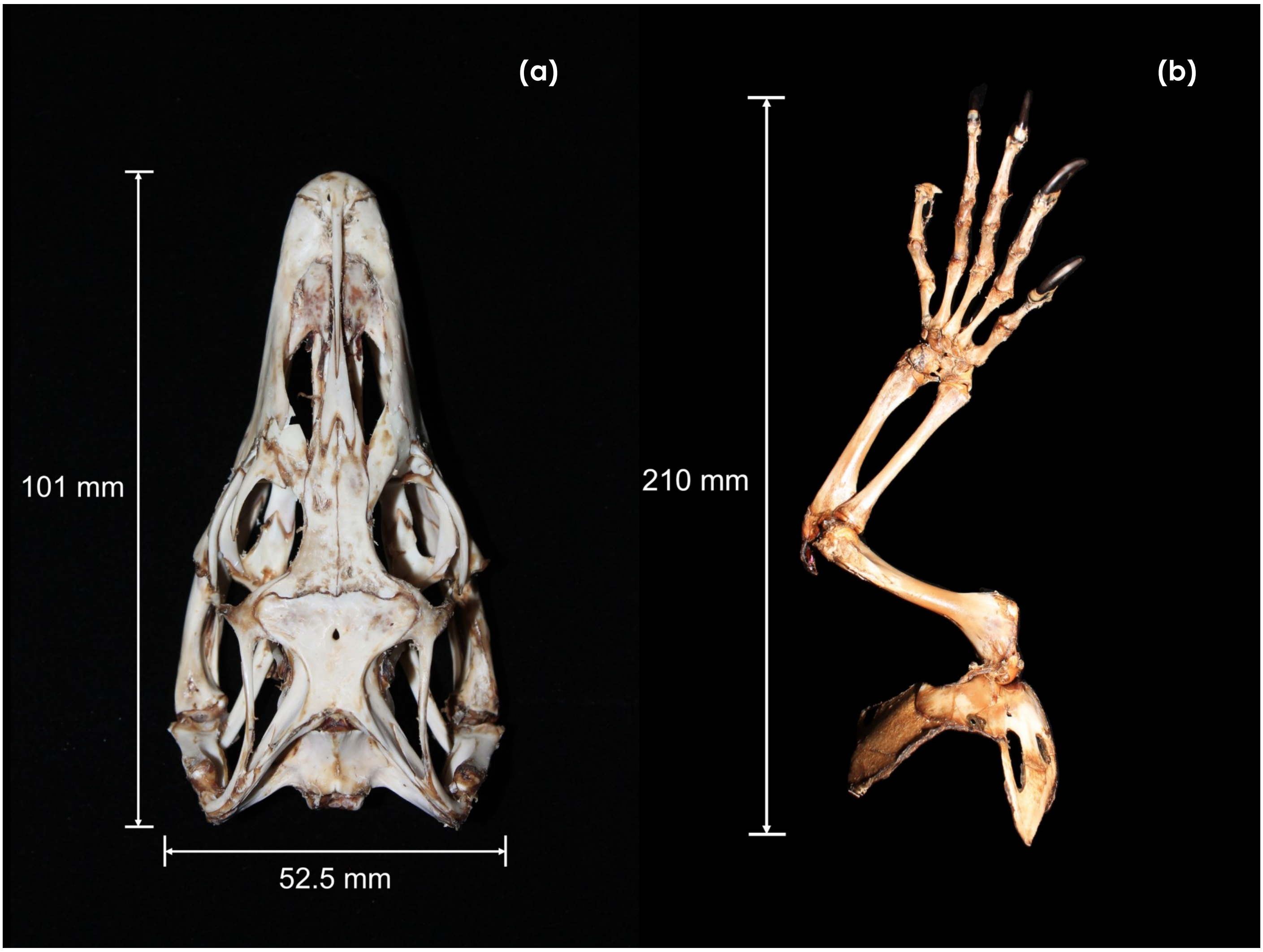


Figure 1.: Varanid (a) skull and (b) front left limb, cf. Figure ?? for anatomical name and ID.

**(a)**

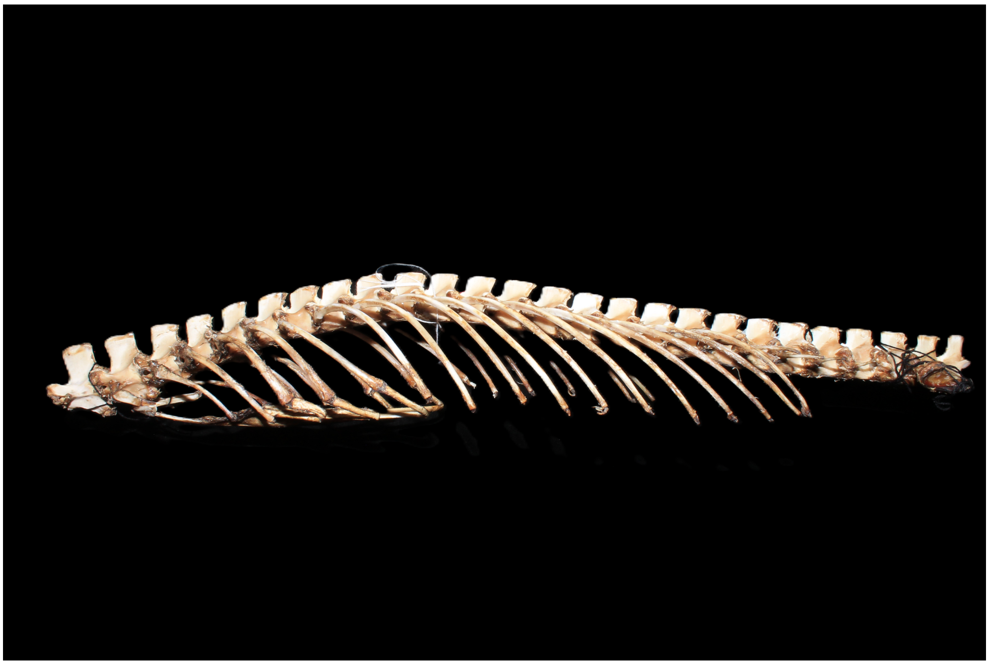

**(b)**

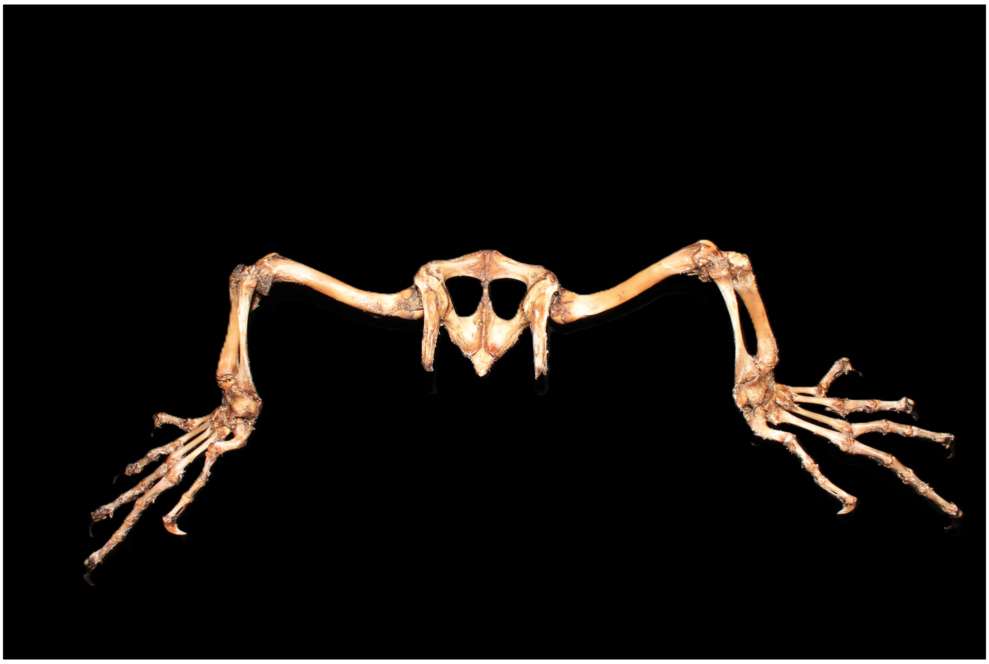

Figure 2.: Images showing (a) lateral view of spine (ID 3.0) and (b) dorsal view of hind limbs and pubic bone (ID 6.0-7.0-8.0).

**(a)**

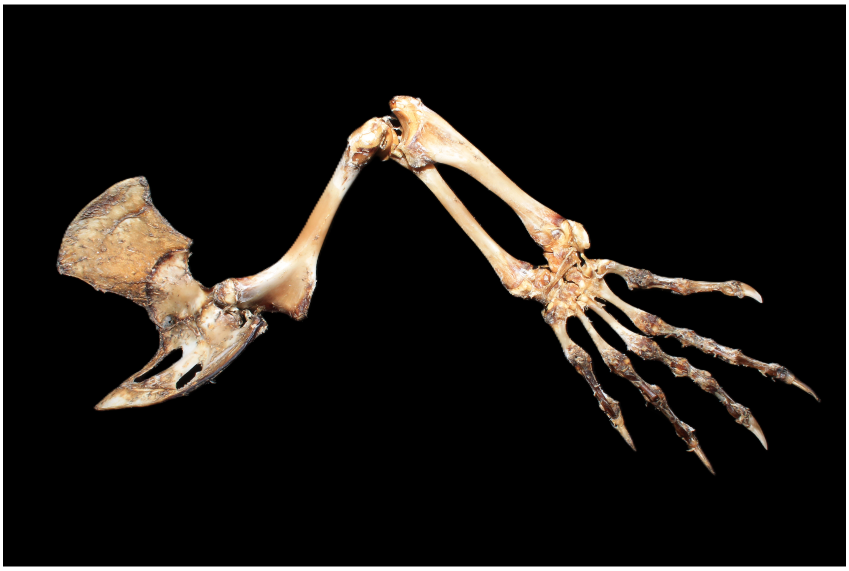

**(b)**

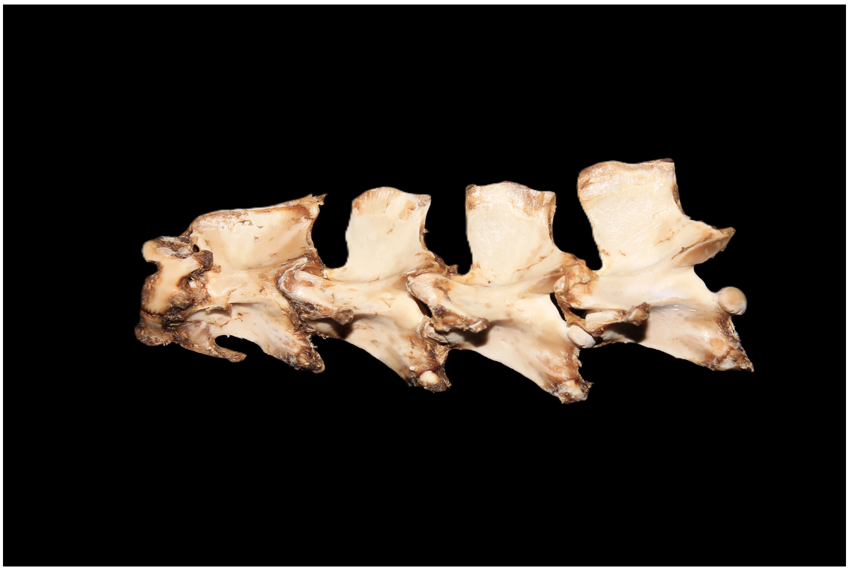

Figure 3.: Images showing (a) ventral view of right forelimb (ID 5.0) and (b) lateral view of neck bones (ID 2.0).

**(a)**

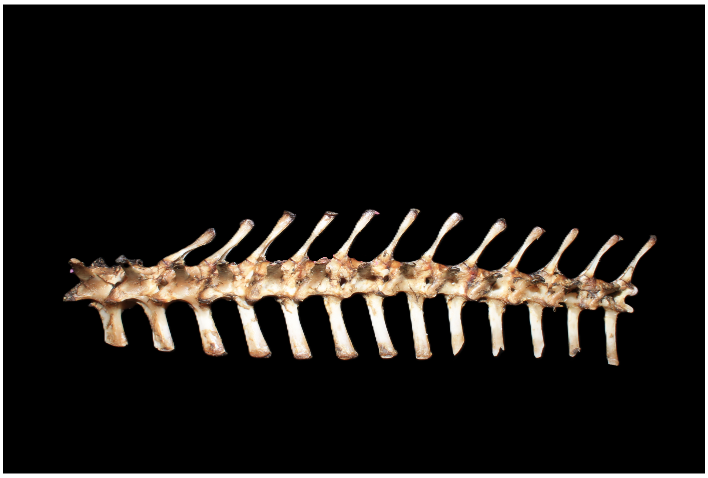

**(b)**

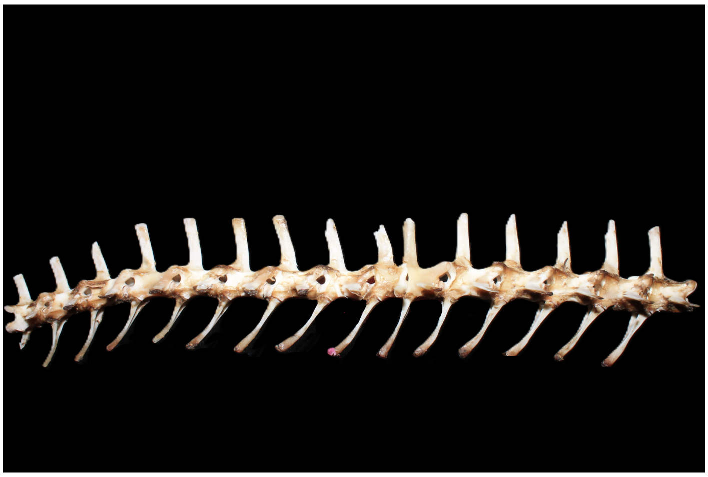

**(c)**

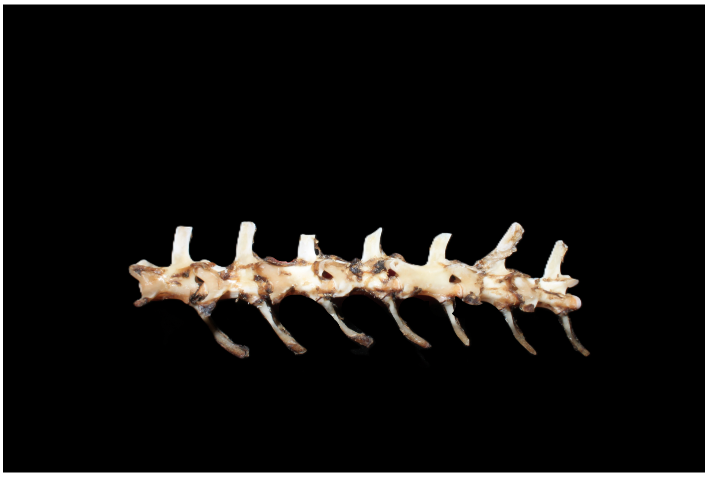

Figure 4.: Images of tail bones: (a) ID 3.4, (b) ID 3.5, and (c) ID 3.6.

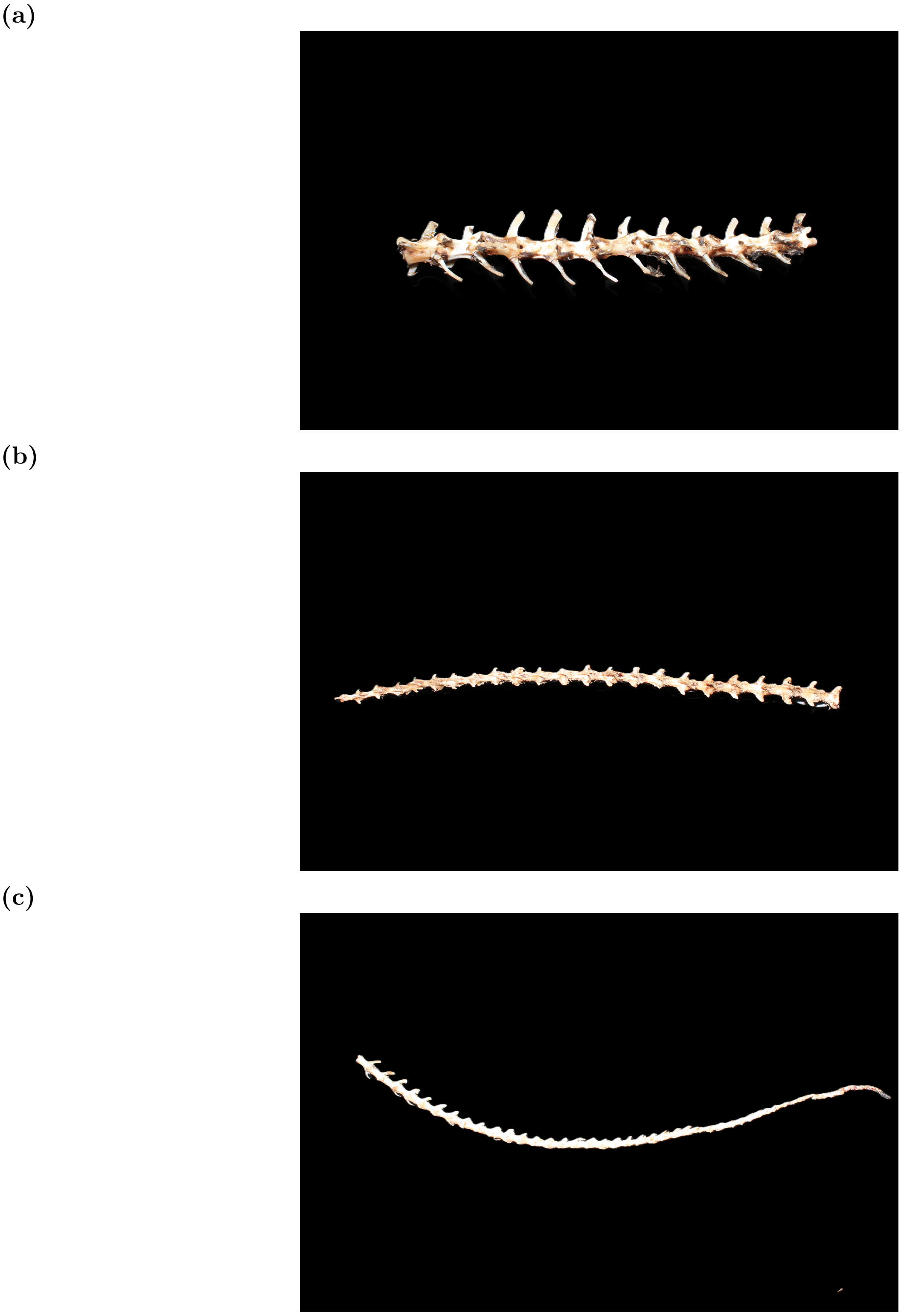


Figure 5.: Images of tail bones: (a) ID 3.7, (b) ID 3.8, and (c) ID 3.9.

#### 2. Skeleton section masses

Table 1.: Masses of intact skeleton sections from first trip to Indonesia

| Bone ID | Common name | Mass (g) |
|---|---|---|
| 1.0 | Skull | 22.7 |
| 2.0 | Neck | 12.6 |
| 3.0 | Spine | 115.1 |
| 3.4 | Tail | 21.4 |
| 3.5 | Tail | 11.9 |
| 3.6 | Tail | 3.9 |
| 3.7 | Tail | 3.9 |
| 3.8 | Tail | 3.9 |
| 3.9 | Tail | 1.5 |
| 4.0 | Left forelimb | 24.7 |
| 5.0 | Right forelimb | 24.4 |
| 6.0 | Pubic bone | 10.4 |
| 7.0 | Left hindlimb | 30.5 |
| 8.0 | Right hindlimb | 31.2 |

#### 3. Lateral trajectory plots for necrobot and varanid

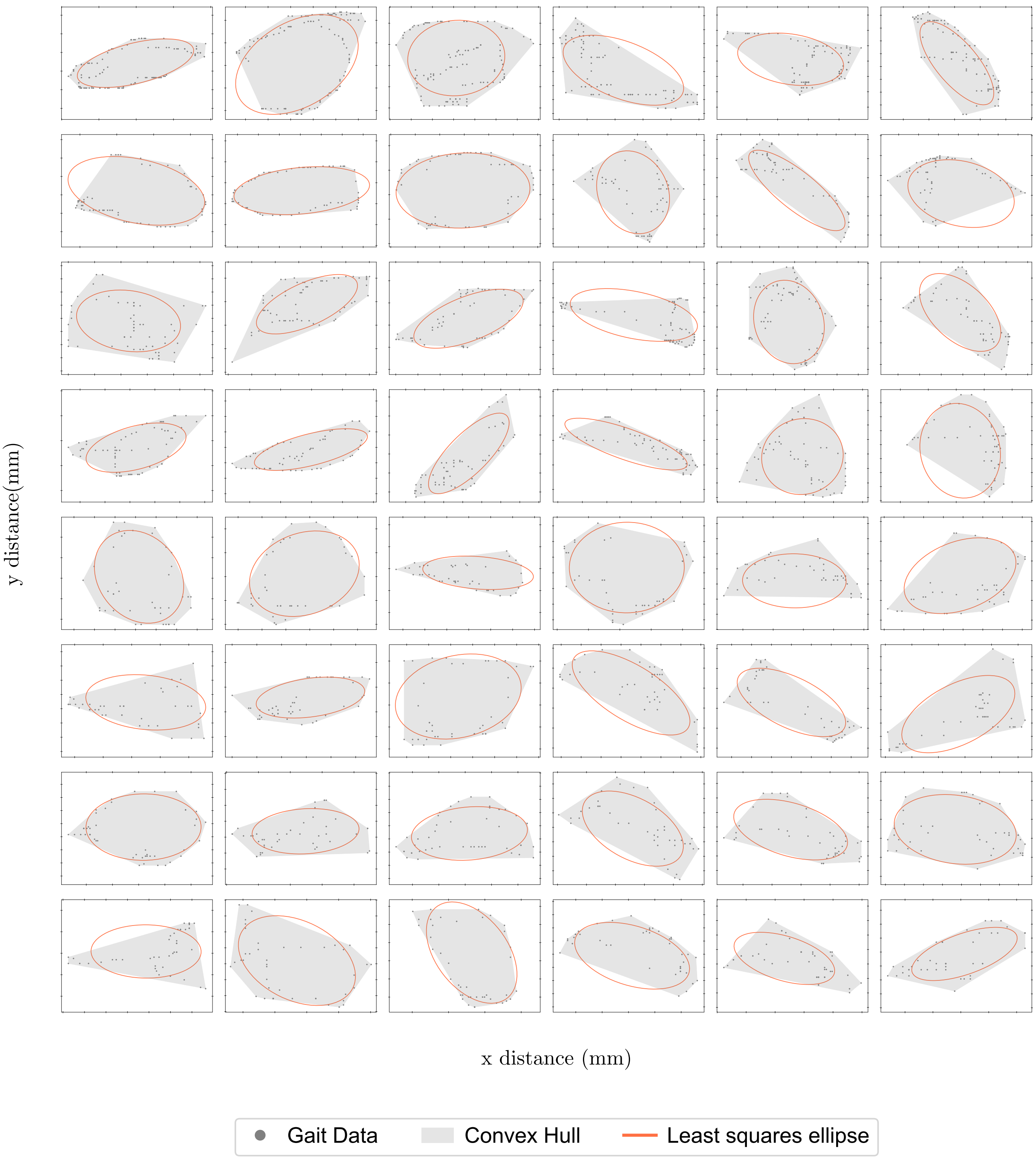


Figure 6.: Necrobot gait lateral ellipse plots for varying speed configurations and leg types

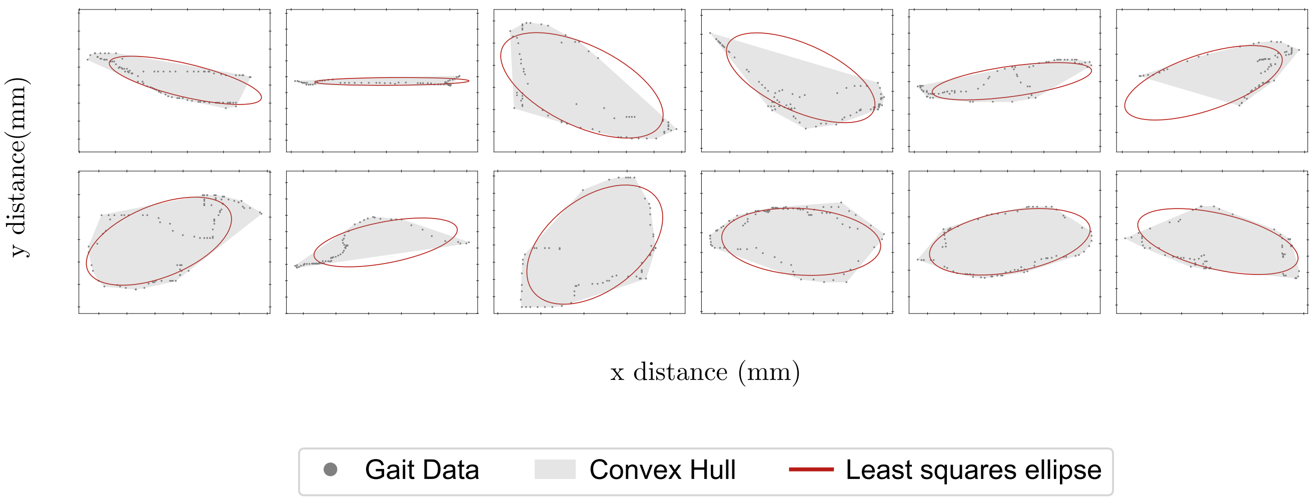


Figure 7.: Varanid gait ellipse plots for different leg positions

#### 4. Illustration of convex hull normalisation

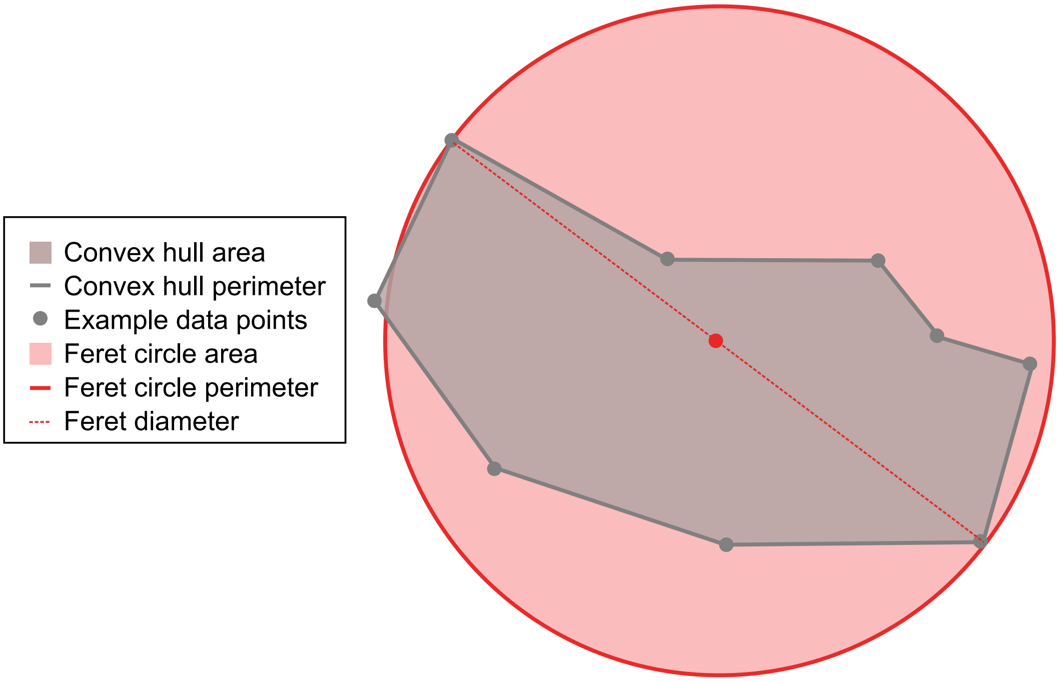


Figure 8.: Example illustration of the convex hull normalisation by the Feret diameter circle

#### 5. Illustration of scatter distance calculations from the fitted curve

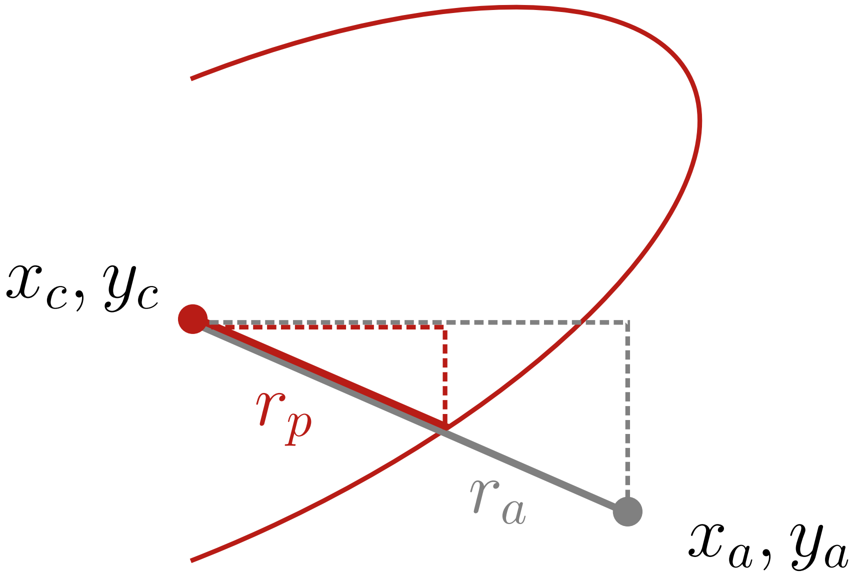


Figure 9.: Illustration of radial distances of actual and predicted points from the centre of the fitted ellipse

#### 6. Comparison of necrobot lateral metrics over different speeds: all plots

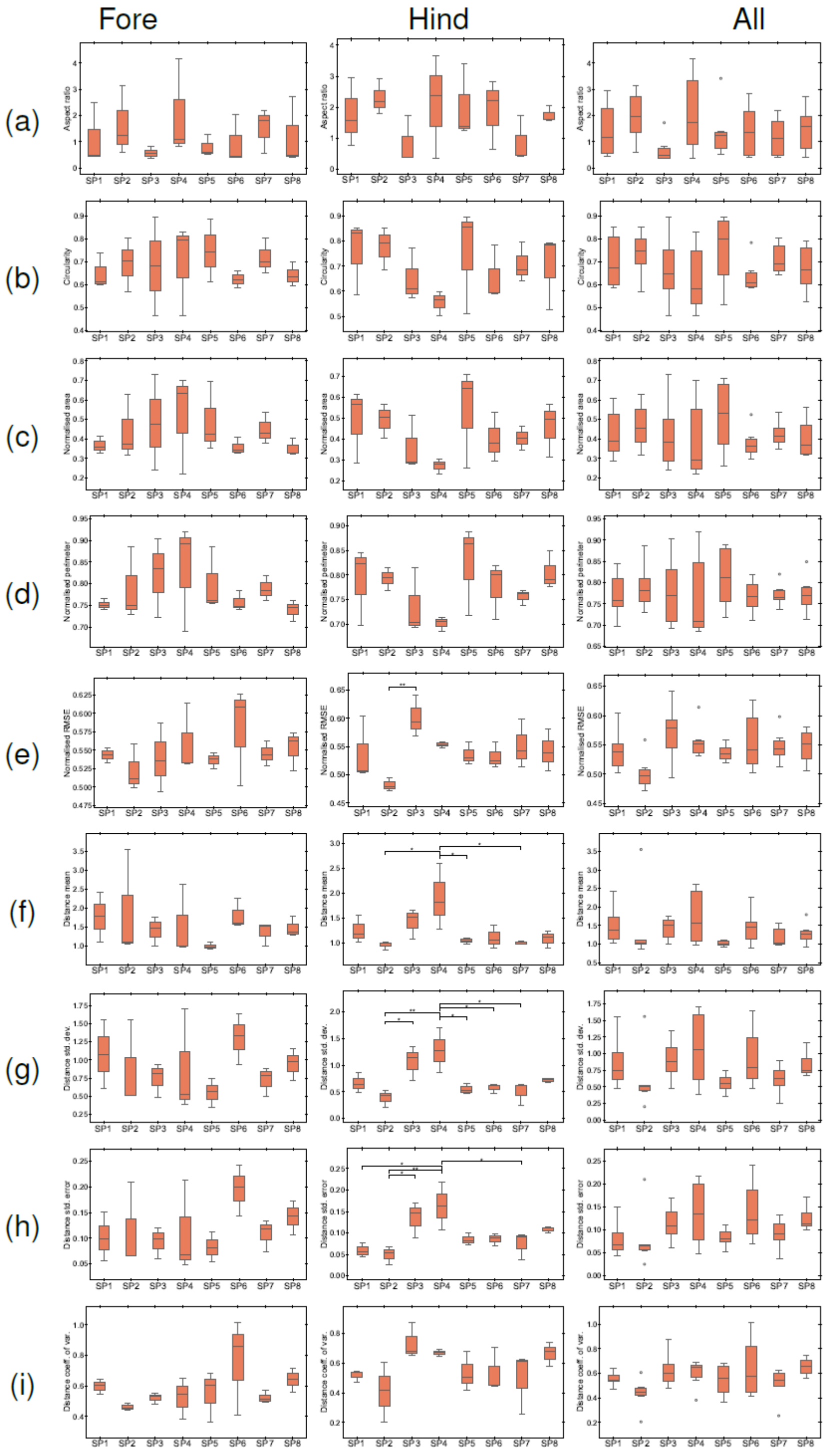


Figure 10.: Box and whisker plots for necrobot speed comparisons for fore, hind and all limbs for (a) aspect ratio, (b) circularity, (c) normalised area, (d) normalised perimeter, (e) normalised RMSE, (f) distance mean, (g) distance std. dev., (h) distance std. error, (i) distance coeff. of var. Boxes indicate interquartile range (IQR); whiskers extend to 1.5×IQR. Significance levels: * p≤0.05, ** p≤0.01, *** p≤0.001; absence of asterisks indicates no significant difference (NS).

#### 7. Comparison of prototype and necrobot lateral metrics against varanid lateral metrics at different speeds: all plots

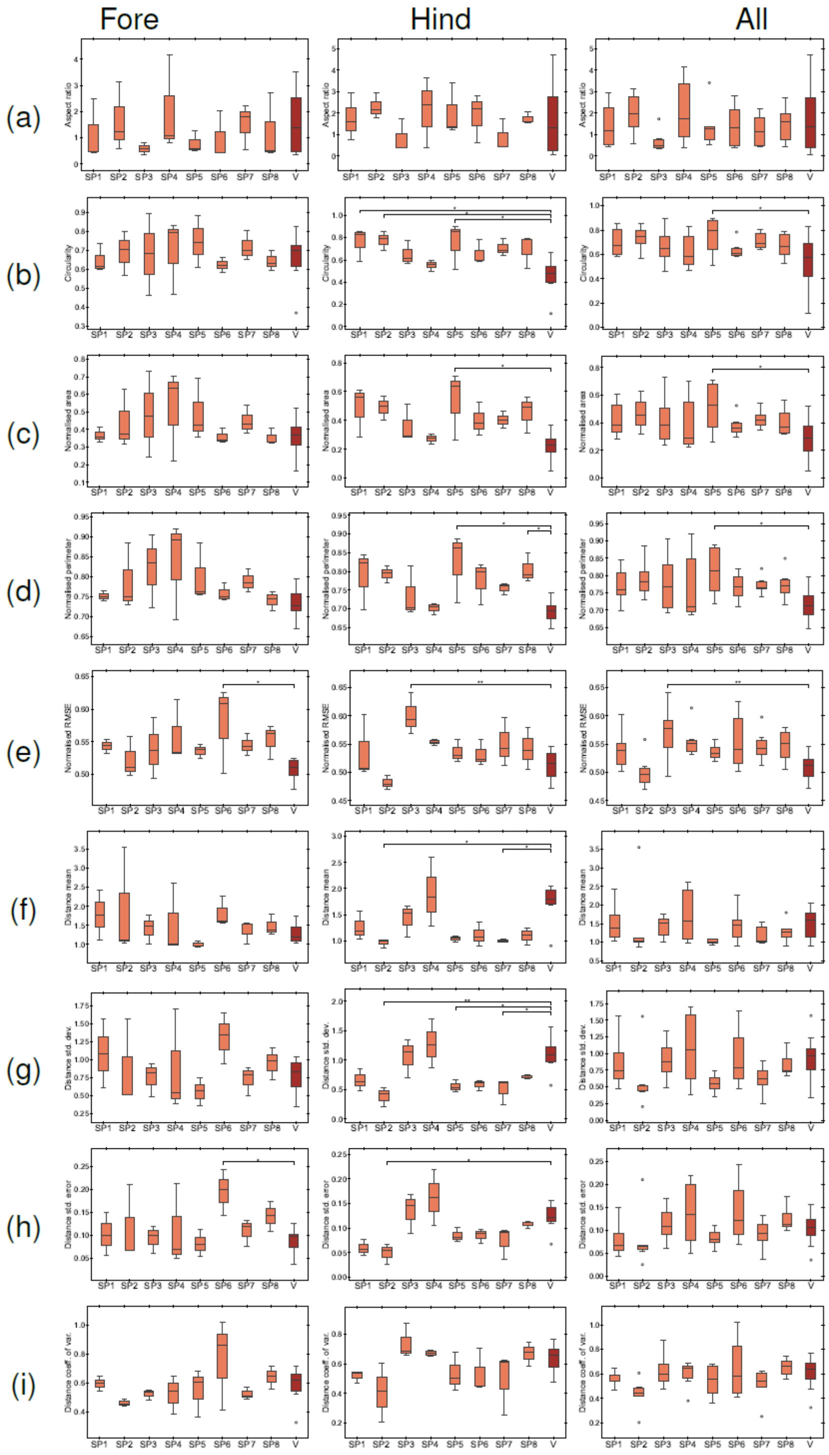


Figure 11.: Box and whisker plots for necrobot metric compared to the varanid for each speed for fore, hind and all limbs for (a) aspect ratio, (b) circularity, (c) normalised area, (d) normalised perimeter, (e) normalised RMSE, (f) distance mean, (g) distance std. dev., (h) distance std. error, (i) distance coeff. of var. Boxes indicate interquartile range (IQR); whiskers extend to 1.5×IQR. Significance levels: * p≤0.05, ** p≤0.01, *** p≤0.001; absence of asterisks indicates no significant difference (NS).

#### 8. Angular and differential curvatures

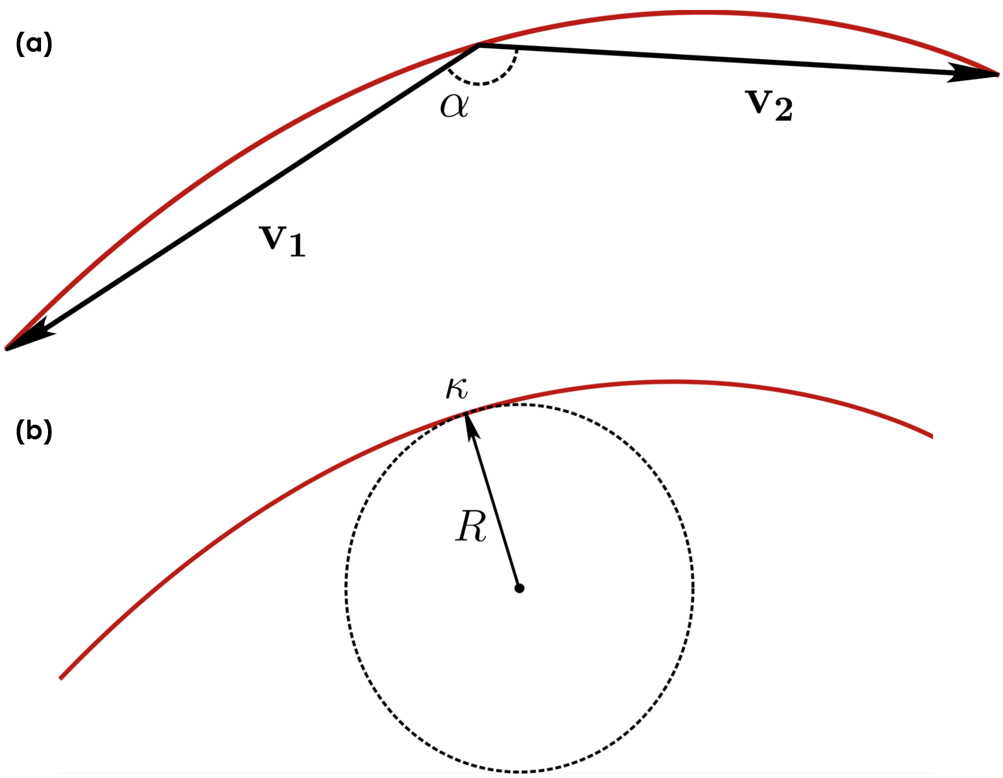


Figure 12.: Illustrations of the fitted polynomial with midpoint curvature calculated by (a) angular curvature using vectors, and (b) differential curvature the fitted polynomial curve.

#### 9. Dorsal spine bending all plots

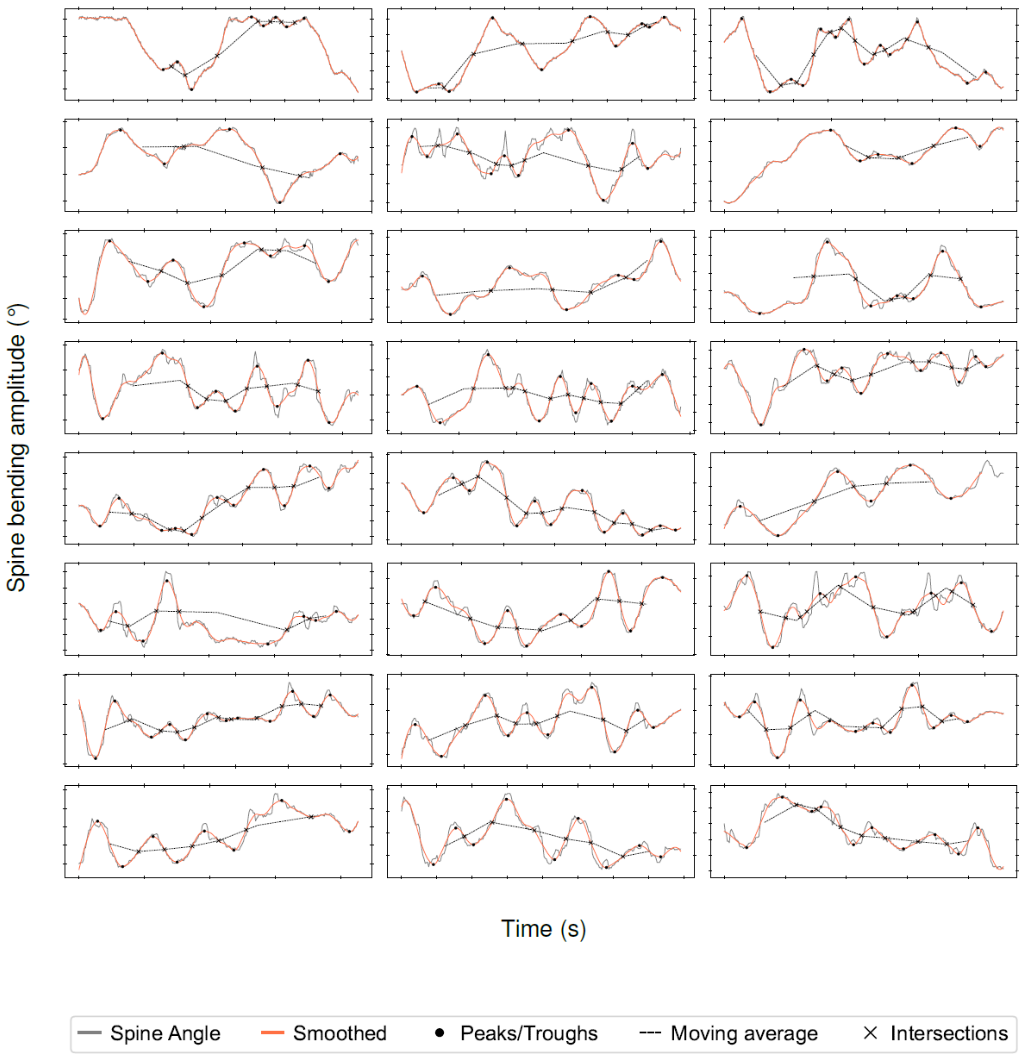


Figure 13.: Necrobot spine bending amplitude at different speeds.

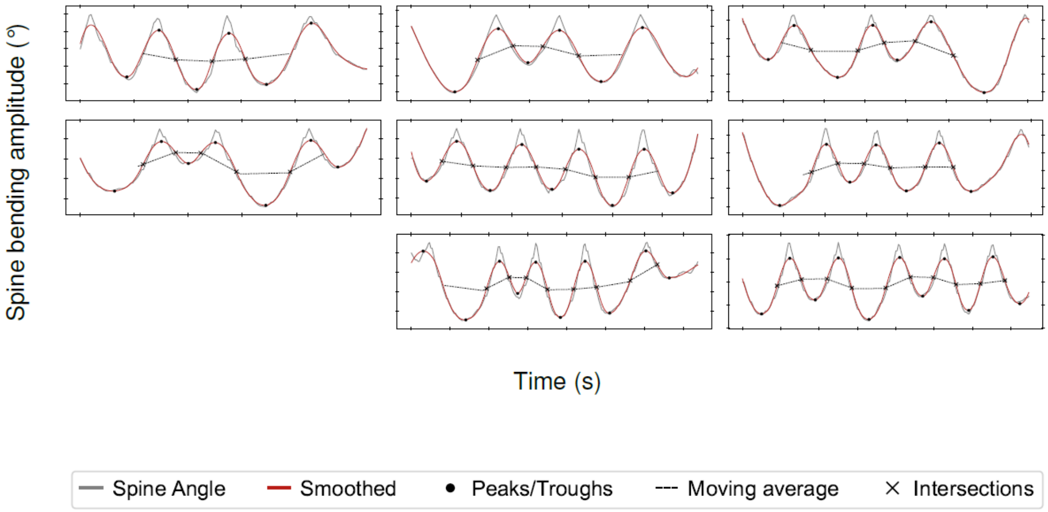


Figure 14.: Varanid spine bending amplitude at different speeds.

#### 10. Dorsal limb sweep all plots

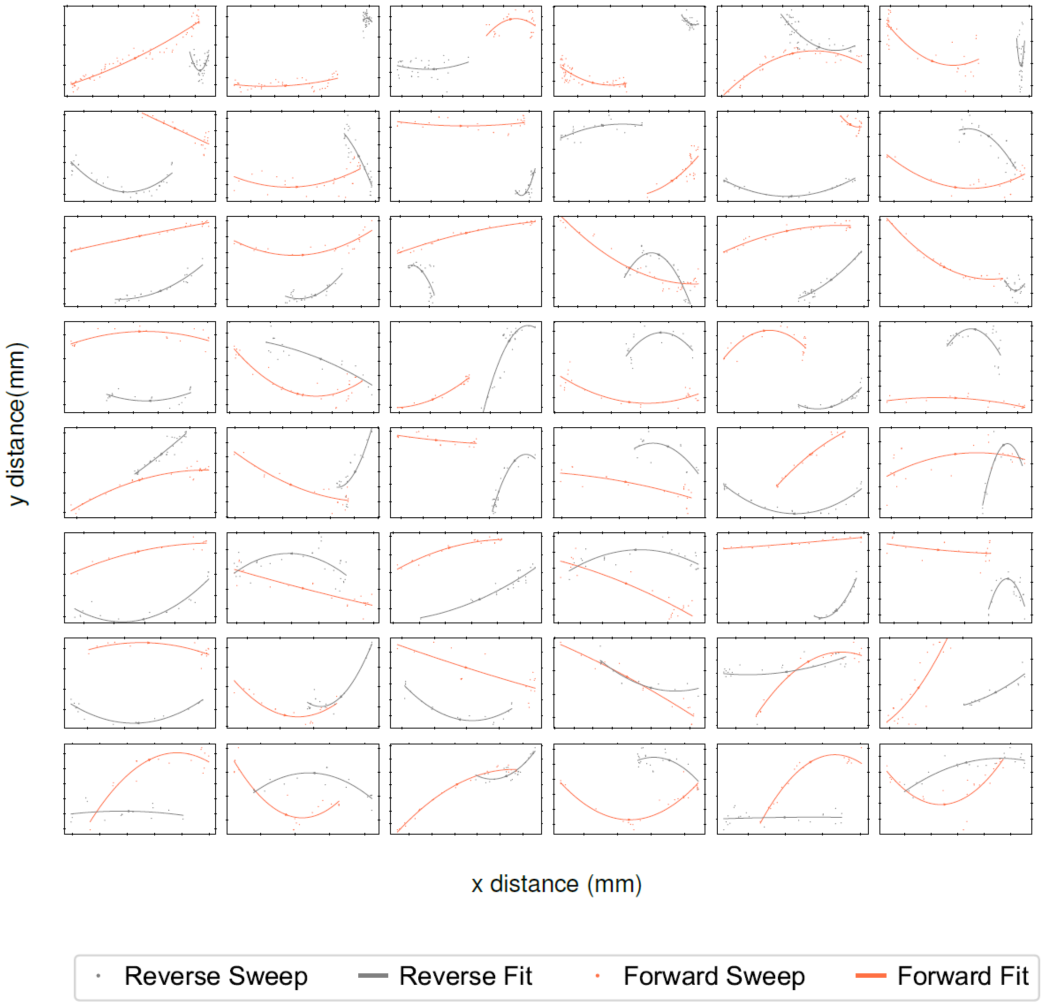


Figure 15.: Necrobot leg sweep paths at different speeds.

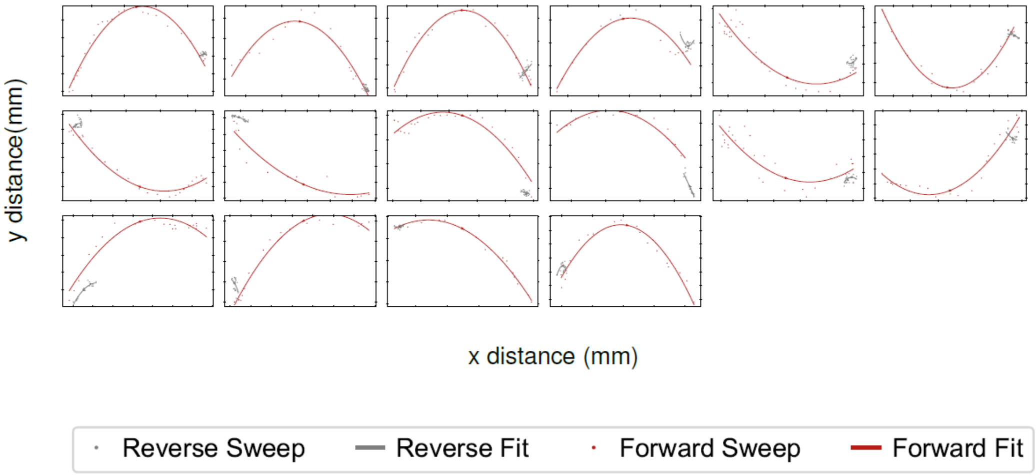


Figure 16.: Varanid leg sweep paths.

#### 11. Dorsal maximum trackway width trace all plots

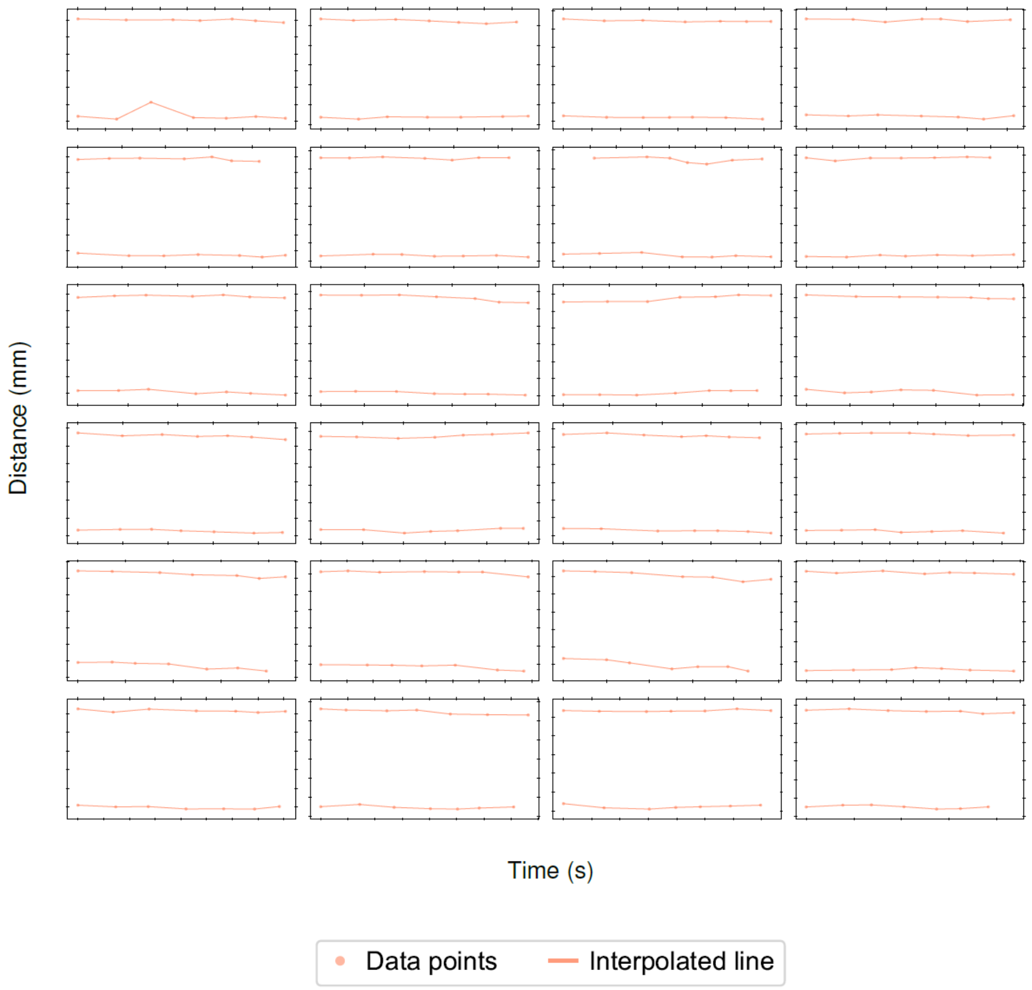


Figure 17.: Necrobot trackway width plots at different speeds.

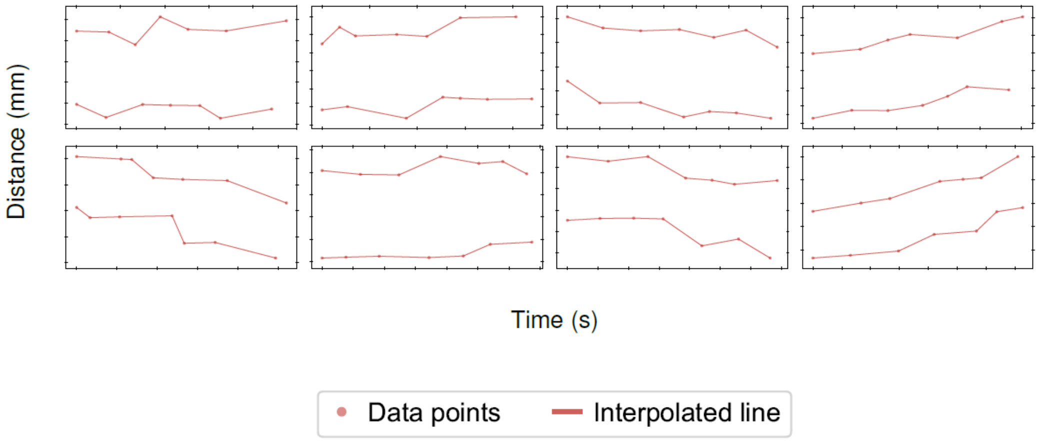


Figure 18.: Varanid trackway width plots.
